# Whole genome regulatory effect of *MoISW2* and consequences for the evolution of the rice plant pathogenic fungus *Magnaporthe oryzae*

**DOI:** 10.1101/2022.02.27.481874

**Authors:** Mengtian Pei, Yakubu Saddeeq Abubakar, Hina Ali, Lianyu Lin, Xianying Dou, Guodong Lu, Zonghua Wang, Stefan Olsson, Ya Li

## Abstract

Isw2 proteins are conserved in eukaryotes and are known to bind to DNA and dynamically influence local chromosome condensation close to their DNA binding site in an ATP-dependent manner making genes close to the binding sites more accessible for transcription and repression. A putative *MoISW2* gene was deleted with large effects on plant pathogenicity as a result. The gene was complemented and a ChIP-sec was performed to identify binding sites. RNAsec showed effects on the overall regulation of genes along the chromosomes for mutant and background strains and this was compared with RNAseq from 55 downloaded RNA-seq datasets from the same strain and found similar. MoIsw2 binding and activities create genomic regions affected by MoIsw2 with high gene expression variability close to the MoIsw2 binding sites while surrounding regions have lower gene expression variability. The genes affected by the MoIsw2 activity are niche-determinant genes (secreted proteins, secondary metabolites and stress-coping genes) and avirulence genes. We further show that MoIsw2 binding sites with the DNA binding motifs coincide with known transposable elements (TE) making it likely that TE-transposition at the binding sites can affect the transcription profile of *M. oryze* in a strain-specific manner. We conclude that MoIsw2 is a likely candidate for a master regulator, regulating the dynamic balance between biomass growth genes (like housekeeping genes) and nich-determinant genes important for ecological fitness. Stress-induced TE transposition is together with MoIsw2 activity a likely mechanism creating more mutations and faster evolution of the niche-determinant genes than for housekeeping genes.

## Introduction

The eukaryotic genome is organized into condensed nucleosomes, limiting access to DNA for transcription factors and more loosely packed regions that are easier to access for interactions with DNA for transcription factors and repressors. The cellular inheritance of this pattern is the basis of animal cell line specialization during embryogenesis (Serrano et al. 2013). Thus, DNA accessibility in fungi also plays an essential role in determining which genes can be efficiently transcribed (Huang et al. 2021).

Imitation Switch 2 (ISW2) belongs to the Imitation SWItch) subfamily (Uniprot) of the SNF2 (Sucrose Non-Fermentable)/RAD54 helicase family. ISW2 proteins contain a Myb/Sant domain close to their C-termini that binds DNA. A role as a transcription factor is the main role often implicated for the Myb-domain containing family proteins in eukaryotes and especially in plants (Wieser and Adams 1995; Mateos et al. 2006; Du et al. 2009; Dubos et al. 2010; Lin et al. 2011; Lin et al. 2012; Prouse and Campbell 2012; Ambawat et al. 2013; Pattabiraman and Gonda 2013; Kim et al. 2014; Baldoni et al. 2015; Dong et al. 2015; Liu et al. 2015; Roy 2016; Matheis et al. 2017; Valsecchi et al. 2017; Verma et al. 2017; Allan and Espley 2018; Mitra 2018; Wang et al. 2018; J. Li et al. 2019; Ma and Constabel 2019; Cao et al. 2020; Li et al. 2024).

A true Isw2 should be involved in gene regulation by regulating the access to DNA for transcription factors and repressors through binding to DNA and controlling nucleosome positioning instead of being a classic transcription factor (TF) (Kagalwala et al. 2004; Fazzio et al. 2005; Hada et al. 2019). Generally, Isw2 proteins have a histone binding domain that interacts with histone 4 of a nucleosome and a catalytic ATPase domain close to the N-terminus that reacts with ATP and changes the Isw2 protein conformation (Whitehouse et al. 2007; Hota and Bartholomew 2011; Dang et al. 2014) so that the nucleosome moves towards the Isw2 DNA binding site (Fazzio et al. 2005), and in that way, transcription is regulated by changing TF and repressor proteins access to DNA. Isw2 is thus involved in targeted nucleosome positioning around these sites. As a consequence of this targeted positioning, a localized nucleosome condensation is created that negatively affects the regulation of the genes at the DNA binding site (Whitehouse et al. 2007) but favours regulation a bit further away from it (Fazzio et al. 2005; Donovan et al. 2021).

There has been much research into Isw2-like protein binding, but it has become clear that *in vitro* mapping of Isw2 binding and nucleosome positioning does not reflect what mechanistically occurs *in vivo* (Fazzio et al. 2005; Donovan et al. 2021), or the very transient nature of the interaction with His4 (Erdel et al. 2010; Erdel and Rippe 2011). The interaction is transient in that the nucleosomes closest to the Isw2 binding sites get dynamically positioned by very frequent (seconds) interaction with the Isw2 protein in an ATP-dependent manner. The activity keeps the closest nucleosome(s) almost immobile, while nucleosomes further away get more freedom to move and genes there are easier to access and regulate. (Donovan et al. 2021). Experiments have shown that both the largest subunits of the Isw2 complex, Isw2, and Itc1, are needed for robust, target-specific binding to DNA while Isw2 alone is sufficient for basal-level binding (Fazzio et al. 2005).

Furthermore, Isw2 is known to bind DNA preferentially at intergenic regions (Whitehouse et al. 2007) where transposable elements (TEs) are commonly located (TE target sites). These sites are staggered cut palindromic target sites (Linheiro and Bergman 2008). In addition, TEs are involved in stress adaptation and host specialization in *Verticillium dahliae* (Faino et al. 2016) as well as in the rice blast fungus *Magnaporthe oryzae* studied here (Chadha and Sharma 2014; Yoshida et al. 2016) and should especially be needed when the host plant detects the fungus as a pathogen at the transition between the endophytic to necrotrophic infection phase of the fungus and thereafter (Cao et al. 2022).

*ISW2* is highly expressed and often considered a “housekeeping” gene stably expressed under normal conditions (Machné and Murray 2012). The stable expression of housekeeping genes has recently been challenged, and many genes traditionally considered stably expressed are unreliable when expressed outside relatively narrow conditions. Thus better methods for selecting housekeeping genes that better fit a specific set of conditions or tissues have been employed (Stanton et al. 2017). On the other hand, the change in ratios of target genes to housekeeping genes reflects the actual change in gene expression ratios of 2 genes of interest. In a previous paper from our lab (Zhang, Zhang, Liu, et al. 2019), we used correlations of expression profiles between genes of interest to probe possible gene functions. That can be done in transcriptomic data from many labs and is especially useful if the calculations are done for all data and not just average value data. Then, also the variation between replicates can be used to calculate the correlation between any two genes’ expressions since these ratios reveal the dependencies of one gene’s expression on the other gene, or both genes’ expressions on a third. Using ratios of gene expressions can thus reveal putative gene functions and help identify genes with specific putative functions that should correlate with the expression of well-known orthologous genes (Zhang, Zhang, Liu, et al. 2019).

In this study we studied the putative *MoISW2* in *M. oryzae* as an *ISW2* gene that affects gene regulation close to the MoIsw2 protein binding sites in the fungal genome. Deletion of the gene negatively affected pathogenicity and stress tolerance. We further show that MoIsw2 binding is at TEs within the genome and that genes close to the binding sites are more differentially regulated (up or down). The affected genes are mainly niche-determinant genes; this, combined with TE transpositions, is a likely mechanism for biased evolution (Monroe et al. 2022) in *M. oryzae* with a faster mutation rate for affected genes.

## Results

### Bioinformatics similarity and domain structure

The protein encoded by the MGG_01012 gene is annotated as a putative MoIsw2 in the NCBI database. From its N to C termini, Isw2 proteins consist of DExx, Helicase C, HAND, SANT and SLIDE domains (Hopfner et al. 2012) (Supplementary method 1 for further comparisons). Thus, the MoIsw2 protein is most probably an Isw2 protein of the IswI and Isw2 type (**Fig 1A)**

**Figure 1.**
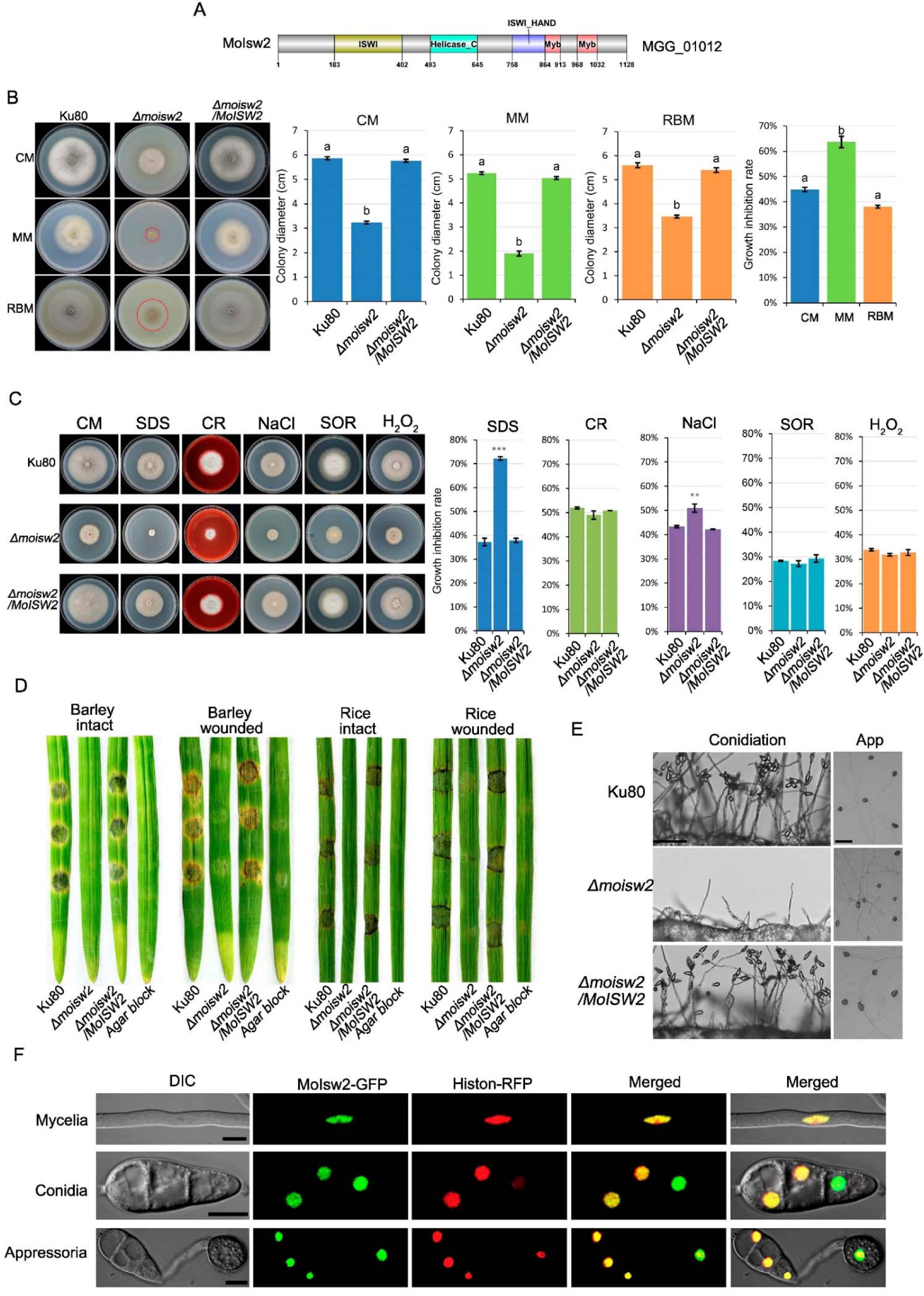
Domain structure, knockout phenotypes compared to the background Ku80, and localization of MoIsw2. (A) Domain structure (B) Growth phenotypes and growth on complete medium (CM) minimal medium (MM) and rice bran medium (RBM). (C) Stress phenotypes on complete medium (CM) with added Sodium Dodecyl Sulfate (SDS), Congo Red (CR), salt (NaCl), sorbitol (SOR) or hydrogen peroxide (H_2_O_2_). (D) Infection phenotypes on barley and rice leaves. (E) Conidia formation and appressoria formation from hyphae. (F) Subcellular localization in mycelia, conidia and appressoria MoIsw2-GFP localizes to nuclei (Histon1-RFP nuclear marker). The same letter on the bars indicates P_same_ > *0.05. The stars above bars indicate significant differences from the Ku80 controls ** P_same_< 0.01 ***P_same_<0.001.

### Deletion phenotypes of *ΔMoisw2* mutant and subcellular localization of the MoIsw2-GFP protein

The *MoISW2* gene was deleted in the *M. oryzae* Ku80 background strain and the deletion was confirmed by Southern blot (**Fig S1**). The effect of this mutation on the vegetative growth of the fungus was tested on three types of media namely complete medium (CM), minimal medium (MM) and rice bran medium (RBM), after incubation at 28 ℃ for ten days. The assays showed significantly decreased colony size of the mutant on all three media when compared to the background strain Ku80 and the complemented strain (*ΔMoisw2*/*MoISW2*) (**Fig. 1B**).

A standard panel of stress treatments added to the CM medium was used to test whether the deletion of the *MoISW2* gene affected the fungal growth rates on different stressing media (Li et al. 2014). The CM medium without the addition of any stress-inducing agent was used as the control medium. The *ΔMoisw2* mutants are strongly inhibited by both SDS (affects membrane integrity) and NaCl (induces osmotic and ionic stress) compared to the controls (**Fig. 1C)**, suggesting that membrane pumps depending on the membrane potential might not be necessarily upregulated and working well in the mutant (Gostinčar et al. 2011).

Furthermore, we tested the ability of the mutant to produce conidia, which are essential for the spread of rice blast disease from plant to plant in the field. We tested this by checking the abundance of conidiophores (which bear conidia) in the various strains. While the WT and complemented strains produced many conidiophores, each bearing many conidia, the *ΔMoisw2* mutant only produced scanty conidiophores with very few conidia (**Fig. 1E**).

Next, the *ΔMoisw2* mutant was tested for pathogenicity on rice and barley leaves and compared with the Ku80 background strain and *ΔMoisw2*/*MoISW2* complemented strain. Since the *ΔMoisw2* mutant produces too few conidia to be used for inoculation, agar plugs were used to infect healthy rice and barley leaves in the infection assay. The assay showed no infection for the *ΔMoisw2* mutant strains (**Fig. 1D**). We reasoned that this could be due to an inability of the mutant to penetrate the plant cuticles. Therefore, we inoculated the fungal strains on aseptically wounded leaves and found that the *ΔMoisw2* mutant could not develop any obvious disease symptoms on the leaves, indicating that *MoISW2* is needed for any successful infection of rice or barley leaves **(Fig. 1D**). From hyphae, *M. oryzae* can form appressoria needed for leaf infection. We found however that dark appressoria formation was not affected by the *MoISW2* mutation, confirming that the cause of no infection in the infection assay was probably not a complete lack of appressoria formation (**Fig. 1E**). Finally, the Moisw2/*MoISW2-GFP* complementation was used to visualize the subcellular localization of MoIsw2. This was performed in a strain expressing Histon1-mCherry (RFP) (as a nuclear marker)(Zhang, Zhang, Chen, et al. 2019) because we expected that MoIsw2-GFP should accumulate in the nucleus as its orthologues do in other eucaryotes. The MoIsw2 protein was observed to consistently colocalize with the nuclear marker in hyphae, conidia and appressoria (**Fig. 1F**).

Taken together, these results suggest that MoIsw2 localize to nuclei and directly or indirectly regulates genes encoding proteins that in several ways strengthen the cell wall and cell membrane barrier important for growth inside a plant and might explain why the deletion of the gene decreased the pathogenicity of the fungus.

### Correlation analyses of *ISW2* expression with expressions of nucleosome histones and ITC1

Histone 1 (*HIS1* gene) is involved in epigenetic activities by changing access to DNA, and its expression is linked to silencing activity (Willcockson et al. 2021). The active Isw2 protein complex containing Itc1 binds DNA and Histone 4 (Donovan et al. 2021) also affects access to DNA. We used downloaded published expression data from the course of rice leaf infection from many experiments for the strain we use to investigate correlations between the putative *MoISW2* and the MoIsw2 complex gene candidates in the same way as we have done before for other genes using the same data (Zhang, Zhang, Liu, et al. 2019). We now investigate if the putative *MoISW2* is transcriptionally coregulated with genes that encode proteins MoIsw2 need to interact with to perform MoIsw2 local site-specific epigenetic activities. The *MoHIS1* expression increases steeply with *MoISW2* gene expression (**Fig. 2**). In *M. oryzae,* there are two genes (MGG_06293 and MGG_01160) *MoHIS4a* and *MoHIS4b* respectively that are predicted (NCBI) to encode an identical His4 protein. Thus the sum of these gene regulations should best reflect the regulation of the MoHis4 protein. Isw2 further works together with ITC1 in a complex that interacts with His4 (Kagalwala et al. 2004). Thus, the expression of these 4 putative genes (*MoISW2*, *MoITC1* and *MoHIS4a*+*MoHIS4b*) producing 3 proteins (MoIsw2, MoItc1 and MoHis4) should be correlated in *M. oryzae,* and they are nicely correlated in the data from plant infection (**Fig. 2** and **Fig S2**). The closest correlation of *MoISW2* expression is even with *MoHIS4a*+*MoHIS4b.* Thus, it points to that the putative MoIsw2 is a real Isw2 as these proteins encoded by these genes are known to interact with His1, Itc1 and His4 in a complex (Fazzio et al. 2005). Furthermore, the expression of *MoHIS2B* is related to the overall growth rate (and thus *de novo* DNA synthesis rate) (Zhang, Zhang, Liu, et al. 2019) and is correlated with *MoISW2* but not with a steep slope. This points to that a smaller amount of MoHis2B is needed than His4 for handling the de novo DNA synthesis per cell than what is needed when Isw2 activity is high. That in turn could mean a lowering of the DNA-synthesis rate with higher *MoISW2* expression. Also, the expression of *MoHIS3* but not *MoHIS2A* is significantly correlated with the putative *MoISW2*. **(Fig. S3).** Although the MoIsw2 protein interacts directly with His4 according to the literature such interactions could not be detected using a yeast two-hybrid assay (data not shown) and the lack of positive results in this assay is likely due to the very transient nature of the interaction (Erdel et al. 2010; Erdel and Rippe 2011).

**Figure 2.**
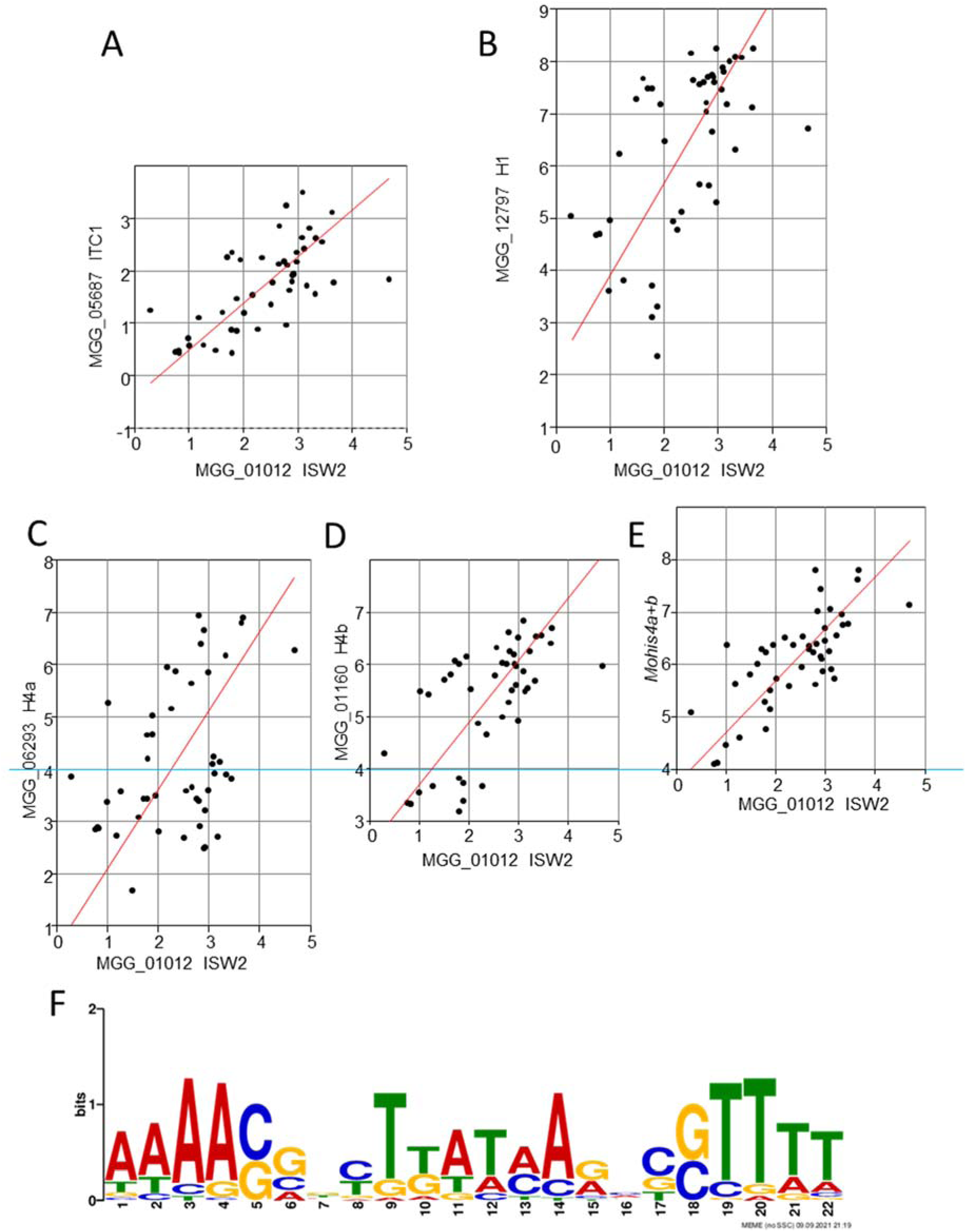
Log2 Reduced Major Axis (RMA) correlations of a putative *ITC1*, the linker histone *HIS1* and the two putative *HIS4* genes putatively expressing the same protein known to interact with the expression of the putative *MoISW2* (x-axis) in published RNAseq data at different stages of plant infection. Each dot represents the two genes’ expression in a separate RNAseq dataset. RMA fitting was used to handle the problem of errors in both X and Y axis values, an inherent consequence of plotting the expression of 2 genes against each other. Each dot corresponds to the value from one transcriptome. (**A**) ITC1, P(uncorrelated)=1.57E-7 (**B**) *HIS1*, P(uncorrelated)=1.48E-5 (**C**) *HIS4a* P(uncorrelated)=0.0044 (**D**) *HIS4b* P(uncorrelated)=3.81E-7 (**E**) *HIS4a* + *4b* since both *HIS4* genes are predicted to encode for the same protein, P(uncorrelated) 5.82E-10. All plots are shown with equal x and y scale gradings so that it is easy to compare slopes between graphs visually. Also, note that figures **C**, **D**, and **E** have been positioned so that Log 4 on the Y-axes are on the same line so that the effect of the addition, Log2(His4a+His4b), can be seen visually. (**F**) A frequently occurring palindromic motif was found in the ChIP-seq data using MoIsw2 as bait. It was found in 196 sequences.

Thus, the putative *MoISW2* gene likely encodes a putative MoIsw2 protein involved in targeted local chromatin compaction working with MoHis1, MoItc1 and MoHis4 (Fazzio et al. 2005; Donovan et al. 2021; Willcockson et al. 2021). 2021). From now on, we skip “putative” as an attribute and investigate if MoIsw2 has the expected functions as an Isw2 and what other effects it has on the biology of the fungus. Furthermore, the local chromatin compaction and the regulatory effect caused by MoIsw2 should be dynamic and stronger the more the gene and the protein it encodes are expressed, and the more ATP there is available for the dynamic interaction.

### ChIP-seq analysis of MoIsw2 finds DNA binding motifs

We made a MoIsw2-GFP fusion protein to perform a ChIP-seq analysis to find conserved DNA binding palindromic motifs for the binding of MoIsw2 to *M. oryzae* DNA sequences.

As mentioned in the introduction, Isw2 proteins are known to preferentially bind DNA at intergenic regions where transposable elements (TEs) are commonly located (Whitehouse et al. 2007) with staggered palindromic motif target sites (Linheiro and Bergman 2008). TEs are also mainly located intergenic in *M. oryzae* in previous unmapped DNA (Chadha and Sharma 2014; Bao et al. 2017) and appear to affect secreted protein expression important for pathogenesis differently in different *M. oryzae* strains (Bao et al. 2017).

Our new data showed hits mainly for intergenic sequences (**Supplemental Data 1**) where the palindromic retrotransposon sequences are preferentially located (Linheiro and Bergman 2008). For these reasons, we looked for palindromic motifs in the ChIP-seq sequences interacting with MoIsw2-GFP. For this analysis we used the MEME website and as input all sequences (peaks) with the ChIP-seq sequences. In addition, we also searched the ChIP-seq sequences for previously identified RT sequences (Bao et al. 2017).

Three palindromic motifs were found to be common (**Supplemental Data 2**) with 47, 196 (**Fig. 2F)**, and 32 occurrences for motifs 1, 2 and 3 respectively, located at 267 DNA locations. The large majority of these locations are intergenic and only a few ChIP-seq sequences had more than one motif hit. We investigated in detail the motif with 196 hits and the majority of these hits were indeed intergenic (113 intergenic locations), while the rest (63) were in the promoter region of a gene. Using a TOMTOM search at the MEME website, we found that the most common palindromic motif 2 is similar to a known human Myb-binding protein containing a DNA binding motif (**Fig. S4, Supplemental Data 3**).

### MoIsw2 DNA binding and its influence on close-by avirulence gene expression

Transposable elements interacting with MoIsw2 may play a direct role in a pathogen-plant arms race if they affect the expression of the nearby avirulence genes (Chadha and Sharma 2014; Bao et al. 2017) through targeted nucleosome condensation (Bourras et al. 2016; Fijarczyk et al. 2022; Feurtey et al. 2023). Avirulence genes are often effectors the pathogen needs to efficiently cause disease and parasitize the host, but different cultivars of the host acquire resistance against these proteins naturally or through human exploitation in plant breeding programs to develop disease-resistant plant varieties (Jones and Dangl 2006). For these reasons, avirulence genes should preferably have become placed adjacent to the MoIsw2 binding site through evolution, and their expression should be affected differently depending on their closeness to the MoIsw2 binding site.

We first made a list of all known avirulence genes noted for *M. oryzae* at NCBI (**Table 1, Supplemental data 9**) to test the above. There are 16 avirulence genes of different types and in addition, the avirulence gene cluster of 12 more genes for a cytochalasan-type compound biosynthesis (Collemare et al. 2008; Song et al. 2015). We then made an RNAseq of the background strain *Ku80* and compared it with the *MoISW2* mutant which also could be compared with the downloaded data. The cytochalasan cluster is specifically activated at early hours post-infection (HPI) during penetration and produces a secondary metabolite recognized by the *R* gene *Pi33* in resistant rice cultivars (Collemare et al. 2008). Two other avirulence genes, Avr-PWL1 and Avr-PWL2 (Dioh et al. 2000), together with the previous, make 18 classic avirulence-type genes, excluding the cytochalasan gene cluster genes (**Table 1**). Most of the genes in the cytochalasan cluster are very closely positioned to MoIsw2 palindromic DNA binding motif sites, and several are differently expressed in strains Guy11 and 98-06 (**Table 1**), which might lead to the production of different final metabolites from the cytochalasan gene cluster even if the different metabolites function as virulence factors.

**Table 1.** Comparison of the position of avirulence genes compared to the position of the MoIsw2 palindromic DNA binding motif sites in strain Guy11, as well as differential regulation between experiments of the avirulence genes in MoISW2 knockout compared to the background, and differential regulation of the same genes during infection of rice in strain Guy11, and strain 98-06 from published data. Blue-marked cells; are genes that MoIsw2 help regulate only in Guy11. Green-marked cells; are genes that are only regulated during infection by 98-06. DelR and VarR contain variation in expression in RNAseq data between mutant and Ku80, and between 55 downloaded RNAseq datasets from different labs, respectively (see text below for explanation of these measures). Several of the listed avirulence genes are not present in any of the two strains and some are only present in one of them.

| ID. | Annotation | Distance from the closest gene to Molsw2 binding site Guy11 | DelR Guy11 | Guy11 expr (X) | 98-06 expr (X) | VarR Guy11 |
| --- | --- | --- | --- | --- | --- | --- |
| MGG_12447 | cytochalasan | 0 | 19.31 | x |  | 48.20 |
| MGG_08386 | cytochalasan | NP |  |  |  |  |
| MGG_08377 | cytochalasan | NP |  |  |  |  |
| MGG_08378 | cytochalasan | 1 | 15.25 | x |  | 15.25 |
| MGG_08380 | cytochalasan | ? |  |  | x |  |
| MGG_08381 | cytochalasan | ? |  |  | x |  |
| MGG_08384 | cytochalasan | 2 | 0.08 | x | x | **29.55** |
| MGG_08389 | cytochalasan | 2 | 2.33 | x | x | **24.59** |
| MGG_08390 | cytochalasan | 1 | 0.00 | x | x | **56.00** |
| MGG_08391 | cytochalasan | ? |  |  | x |  |
| MGG_15927 | cytochalasan | ? |  |  | X |  |
| MGG_15928 | cytochalasan | NP |  |  |  |  |
|  | Avirulence genes |  |  |  |  |  |
| MGG_07199 | ATR13, RxLR effector | 170 | 0.04 | x | x | 7.27 |
| MGG_10556 | Avr_Pii | NP |  |  |  |  |
| MGG_17614 | Avr_Pii | NP |  |  |  |  |
| MGG_13283 | Avr_Pik | NP |  |  |  |  |
| MGG_15972 | Avr_Pik | ? |  |  | X |  |
| MGG_03029 | Avr_Pita1 | 175 | 37.08 |  |  | 19.74 |
| MGG_07038 | Avr_Pita1 | 169 | 2.00 |  |  | 6.25 |
| MGG_09617 | Avr_Pita1 | NP |  |  |  |  |
| MGG_10927 | Avr_Pita1 | 20 | 0.07 |  |  | **24.00** |
| MGG_03808 | Avr_Pita1 like | NP |  |  |  |  |
| MGG_15370 | Avr_Pita1 like | 1 | 0.28 |  |  | **56.00** |
| MGG_17611 | Avr_Pita1 | NP |  |  |  |  |
| MGG_14981 | AVR_PiTA2 | NP |  |  |  |  |
| MGG_15212 | AVR_Pita2 | ? |  |  | X |  |
| MGG_18041 | Avr_Piz-t | NP |  |  |  |  |
| MGG_03685 | Avr-Pi54 | 238 | 14.11 |  |  | 3.86 |
| MGG_13863 | Avr-PWL1 | ? |  |  | X |  |
| MGG_07398 | Avr-PWL2 | NP |  | X |  |  |
| Sum number 30 |  |  |  | 7 | 11 |  |
| Only in one of the strains |  |  |  | 3 | 7 |  |
|  |  |  |  | In both |  |  |
|  |  |  |  | 4 |  |  |

**Table 1.** Comparison of the position of avirulence genes compared to the position of the MoIsw2 palindromic DNA binding motif sites in strain Guy11, as well as differential regulation between experiments of the avirulence genes in MoISW2 knockout compared to the background, and differential regulation of the same genes during infection of rice in strain Guy11, and strain 98-06 from published data.
| ID. | Annotation | Distance from the closest gene to MoIsw2 binding site Guy11 | DelR Guy11 | Guy11 expr (X) | 98-06 expr (X) | VarR Guy11 |
| --- | --- | --- | --- | --- | --- | --- |
| MGG_12447 | cytochalasan | 0 | 19.31 | x |  | 48.20 |
| MGG_08386 | cytochalasan | NP |  |  |  |  |
| MGG_08377 | cytochalasan | NP |  |  |  |  |
| MGG_08378 | cytochalasan | 1 | 15.25 | x |  | 15.25 |
| MGG_08380 | cytochalasan | ? |  |  | x |  |
| MGG_08381 | cytochalasan | ? |  |  | x |  |
| MGG_08384 | cytochalasan | 2 | 0.08 | x | x | **29.55** |
| MGG_08389 | cytochalasan | 2 | 2.33 | x | x | **24.59** |
| MGG_08390 | cytochalasan | 1 | 0.00 | x | x | **56.00** |
| MGG_08391 | cytochalasan | ? |  |  | x |  |
| MGG_15927 | cytochalasan | ? |  |  | X |  |
| MGG_15928 | cytochalasan | NP |  |  |  |  |

Six other avirulence genes are not expressed in either strain (**Table 1**). Of these 6 genes, half of the genes are close to the MoIsw2 binding site, and 3 are further away in Guy11. Two avirulence genes (MGG_17614 and MGG_15611) are in supercontigs with unknown gene order in our data, so it is unsure how far they are from a MoIsw2 binding site.

### TE position and MoIsw2 DNA binding

We searched the short MoISW2 ChIPseq sequences data for described TEs sequences (Bao et al. 2017) and found described TEs in 92.2 per cent of sequences while the genome outside these ChIPseq sequences is almost devoid of TEs (**Supplemental data 5**). We mapped the MoIsw2 binding sites to the genome as well as the position of all found transposable elements (TEs) (**Fig. 3**). The positioning of the MoIsw2 binding sites and the palindromic motifs we found are close to retrotransposons and the differential regulation. The found expression variation can indicate that MoIsw2-specific targeting is instrumental for stabilizing avirulence gene expression. These data and analyses with additional references are available in supplemental data files (**Supplemental data 4 and 5**). In conclusion, MoIsw2 and RT activities could together be instrumental in creating variation in gene expression profiles between fungal strains.

**Figure 3.**
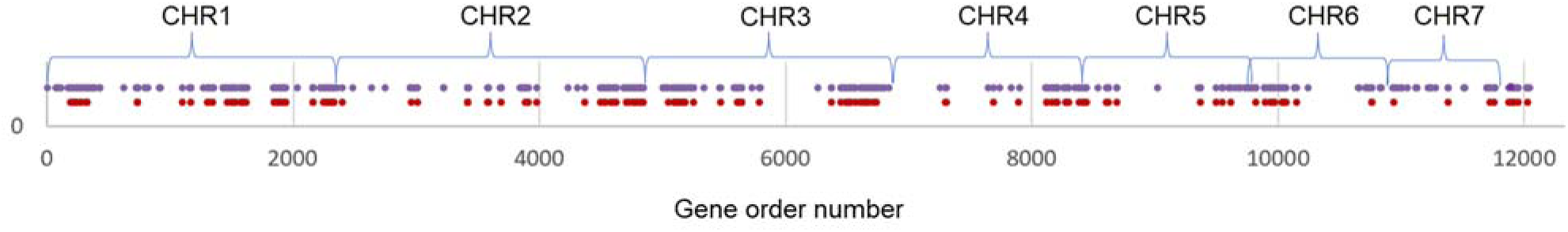
The position of TEs and identified palindromic MoIsw2 binding sites in DNA largely overlap along the whole genome.

### MoIsw2 regulates the genes closest to the MoIsw2 motif sites in the DNA

We next compared the RNAseq for the *Δmoisw2* strain and the background *Ku80* strain for overall gene regulations (**Fig. 4A-B**). There was a general upregulation of the genes closest to the MoIsw2 binding sites in the *ΔMoisw2* mutant (**Fig. 4A**). For these genes, the *ΔMoisw2*/*Ku80* expression ratio is lower when the MoIsw2 is predicted to bind directly in the promoter region than for the closest gene for a predicted intergenic binding of MoIsw2. On the other hand, if absolute (positive or negative) regulation was considered both types of genes, with MoIsw2 binding in the promoter region or intergenic, had similar absolute regulation indicating that many genes with MoIsw2 binding in their promoter region are repressed in *ΔMoisw2*. This suggests that without MoIsw2 activity, repressors can also get access to the DNA (**Fig. 4B**). These observations support MoIsw2 as an Isw2 protein that creates a local nucleosome condensation at specific nucleosomes (Donovan et al. 2021) increasing genes’ either transcriptions or repressions at slightly further distances from the DNA binding sites.

**Figure 4.**
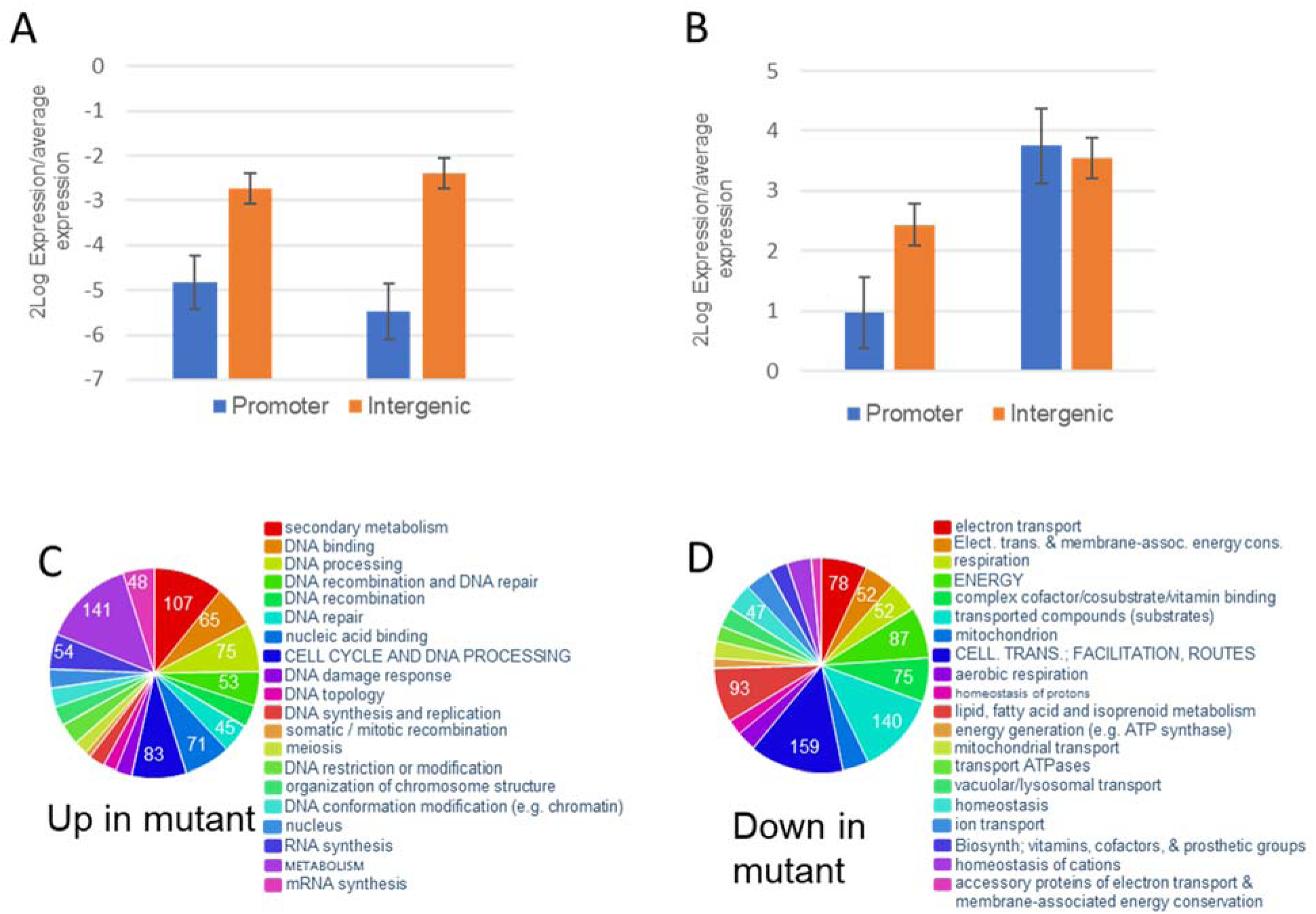
Deletion-induced change in expression of genes next to the predicted palindromic MoIsw2 DNA binding site with hits for the MoIsw2 binding in the ChIP-seq data compared to the average regulation of all genes. (**A**) Expression was, on average higher of genes closest to the binding site in *ΔMoISW2* than Ku80, or with the motif in their promoter region. (**B**) Expression of genes with a binding site in the promoter region and the closest gene when MoIsw2 binding is intergenic. MoIsw2 upregulates both gene types and upregulates genes slightly further away from the binding site, while the absolute (plus or minus) regulation is equal for both types of genes (8-16 times regulation). Error bars indicate SEM (Promoter N=63 Intergenic N=112). (**C**) 20 most significantly enriched upregulated FunCat gene categories in the *ΔMoISW2* compared to the background strain Ku80; Secondary metabolism, DNA-binding, and genes for DNA-related activities and synthesis (anabolism). (**D**) 20 Significantly enriched downregulated genes in the mutant are genes for, mitochondrial activities like electron transport and mitochondrial biosynthesis, respiration, and transport routes (catabolism).

### Overall functional classification of differentially expressed genes in the RNA-seq data

In *ΔMoisw2* grown on MM-medium, 339 genes were significantly upregulated relative to the background *Ku80*. The two largest significantly overrepresented gene categories were secondary metabolism and DNA-binding, while most other categories are related to DNA synthesis, DNA-linked activities and other growth-related processes (anabolism) (**Fig. 4C**). These are the genes MoIsw2 likely negatively regulates in the background *Ku80*. Most downregulated genes in the *ΔMoisw2* are genes involved in mitochondrial electron transport, other mitochondrial processes, transported compounds and other processes connected with uptake and respiration and oxidative phosphorylation (catabolism) (**Fig. 4E**).

In other words, our results indicate that MoIsw2 is involved in regulating the balance between anabolism and catabolism. Between DNA synthesis, which is safest without ATP generation by mitochondrial respiration since that generates Radical Oxygen Species (ROS), and oxidative phosphorylation, which is more effective in generating ATP from the metabolized substrates (Klevecz et al. 2004).

### The local gene regulations close to the MoIsw2 palindromic DNA binding sites fit the Isw2-specific targeted DNA binding model

DNA is wrapped around and slides around nucleosomes of similar sizes, with a constant overall Nucleosome Repeat Length (NRL) (van Holde 1989; Cutter and Hayes 2015; Donovan et al. 2021; Willcockson et al. 2021) and NRLs sizes vary slightly depending on the organism, cell type and cell status (van Holde 1989). To further investigate local regulation around at or close to the binding sites with the found palindromic motif, we investigated our RNAseq data and ordered the MGG-codes for the genes in the order of the genes on the chromosomes (Chromosome order downloaded from BROAD).

Dynamic changes due to chromosome packing affected by MoIsw2 should result in the same regulatory landscape in ween in our experiment and the downloaded data. Since MoIsw2 activity depends on respiratory catabolism activity (nutrient availability in the niche) as mentioned above, the greatest variation in expression between experiments should be for genes positioned close to the MoIsw2 binding sites in the DNA. The change of the measured gene expression responses in our RNAsec data and compared expression responses (positive and negative) in Ku80 (background) strain with the *ΔMoisw2* mutant strain (DelR). We could then test if the variation in expression responses (positive and negative) (VarR) in the downloaded data from 55 RNAsec datasets from several labs shows expression variations at similar positions in the data as were affected by the deletion. We found that the two gene expression measurements were Log-Log correlated (P=5.3E-196) and that the response slope of the correlation was 0.56 with VarR on the x-axis (**Fig. S5, Supplemental Data 11**).

We further calculated the gene distance between each gene and the closest MoIsw2 binding site (**Supplemental Data 10**) and plotted that together with plots of the gene expression change from our experiment (DelR) and the variation in the downloaded data (VarR). In the plot (**Fig. 5**), we also included the positions of the avirulence genes (rings). From these comparisons, one can see that gene regulatory responses along the genome are dependent on MoIsw2 and its binding to DNA. Further, we can identify regions along the DNA with high variation in gene regulation between experiments close (less than 16 genes from the closest motif) to MoIsw2 binding motifs (**Fig. 5** Regions (A-U)) and regions with less variation further away (17-to about 500 genes) from the MoIsw2 binding motifs (**Fig. 5** Regions (1-15)). In **Figure 5** we have marked the positions of the Avir genes as well.

**Figure 5.**
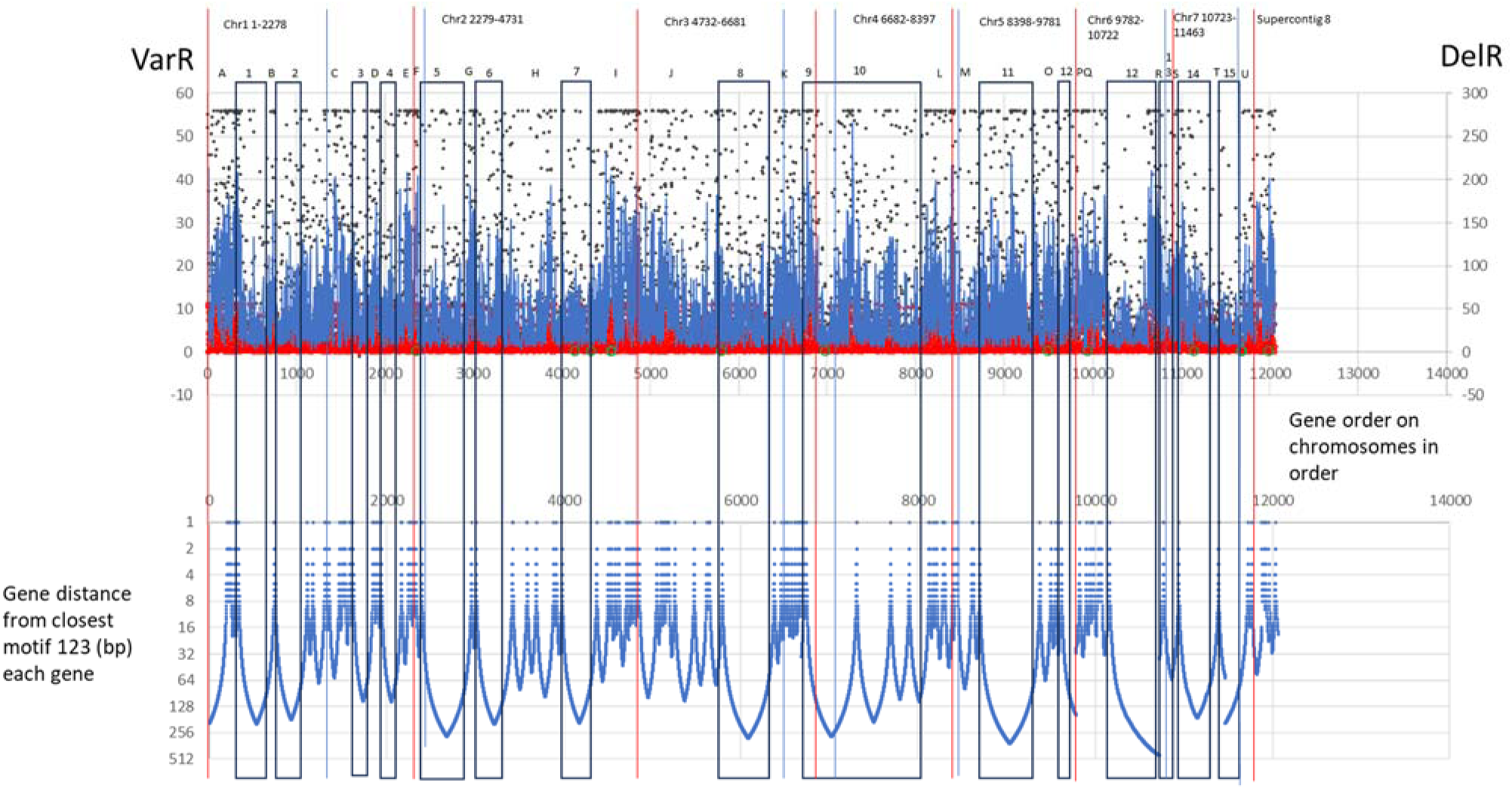
A plot of distances of all genes to the closest motifs of the 123 binding motifs sites (**bottom**) is compared to a plot of the variation of expression across 55 RNAseq experiments (black points and blue 5-point moving average) combined with the difference in gene expression between the *MoISW2* mutant and the Ku80 background (Red points and bright red 5-point moving average (**top**) (DelR). For the VarR data, several genes had at least one condition when their expression was below the detection level while otherwise expressed in these cases the gene expression was set to the lowest detected gene expression. That creates a line of black dots at the top of the figure. The density of the line of dots (top figure) together with areas with less than 16 gene distances from the motif sites paints out the regions of the genome with the most variation in both the VarR and DelR data. The genome contains A-U genomic regions with high variability of gene expression closest (less than 16 genes away) to the MoIsw2 123 DNA binding motifs sites and 1-15 regions with lower variation. Note bottom Y axis is log and reverse, X axis is gene order from 1->12000 on the chromosomes in chromosome order from 1-7 plus supercontig 8 for the whole genome to get the whole genome in order.

For typical housekeeping genes like the genes encoding ribosomal proteins (Zhang, Zhang, Liu, et al. 2019), their expression depends on the overall rate of translation and should be quite constant between experiments. These genes are mainly found further away from the MoIsw2 binding sites and TEs confirming that the ribosomal proteins are in regions with more stable expression (**Fig. 6A**). On the other hand, genes that are important for interaction with the environment as the secreted proteins with identified domains (SP) and smaller other secreted proteins (OSP) (Bao et al. 2017) are positioned in the more variable genomic regions closer to the MoIsw2 binding motifs (**Fig. 6B** and **6C**,

**Figure 6.**
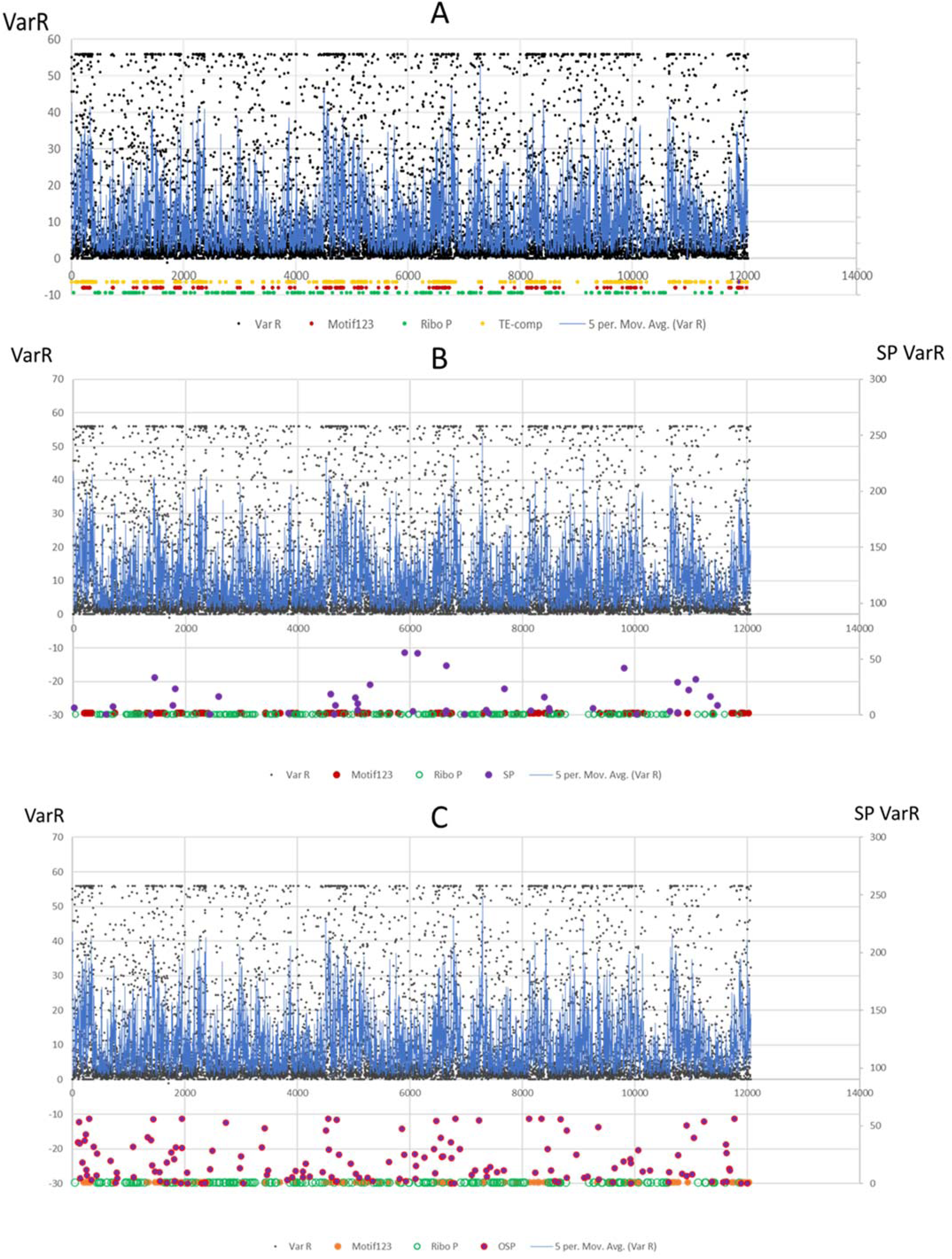
Position of housekeeping ribosomal genes and genes encoding secreted proteins compared to the position of MoIsw2 DNA binding motifs and genes encoding ribosomal genes compared to VarR variability. (**A**) Ribosomal genes. (**B**) Genes for secreted proteins (SP) with conserved domains (generally larger secreted proteins). (**C**) Genes for other secreted proteins (OSP) without any conserved domains identified (generally small secreted proteins and peptides).

### Supplemental Data 6-7)

Other genes that are important in interaction with the environment are secondary metabolites. Instead of searching the literature, we used the AntiSmash web server and submitted the whole DNA sequence from BROAD to get a more extensive list of putative core secondary metabolite genes. We found 64 core secondary metabolite genes and plotted their positions along the ordered gene positions (**Fig. 7A, Supplemental Data 8**) and found that they are mainly positioned close to MoIw2 binding motif sites.

**Figure 7.**
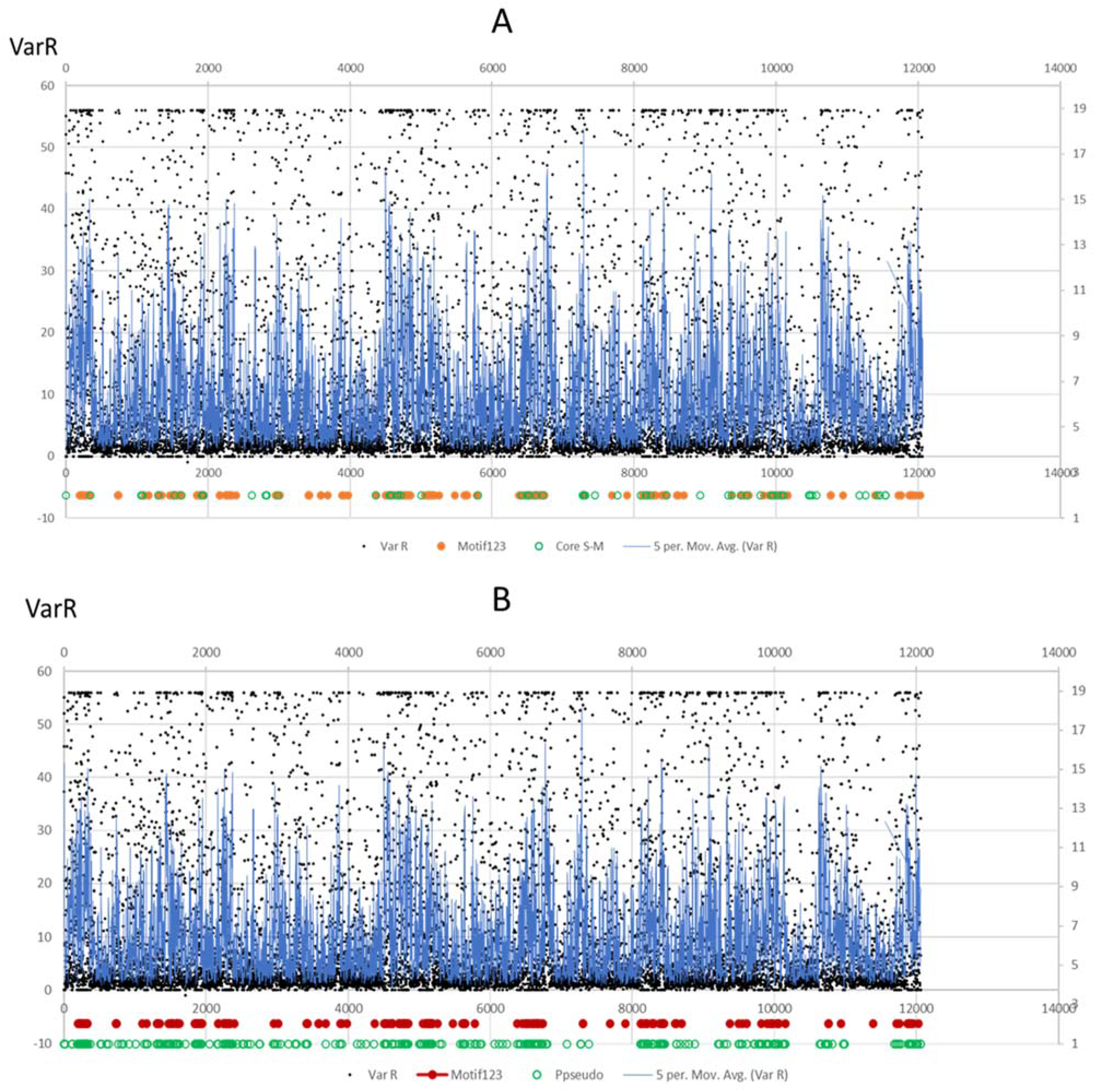
Position of core secondary metabolite genes and potential pseudogenes compared to the position of the MoIsw2 DNA binding motifs and genes encoding ribosomal genes compared to VarR variability. (**A**) Core secondary metabolite genes (green rings). (**B**) Potential pseudogenes not expressed in any treatment (dark green rings). In both A and B, the positions of the 123 MoIsw2 DNA binding motif sites are marked as references.

Epigenetic silencing can if it is persistent from generation to generation result in the silenced genes accumulating deleterious mutations since the genes are never expressed and needed in the ecological niche the organism lives in. We consider genes not expressed in any of the downloaded RNAsec datasets as potential pseudogenes (PP). Since the misuse of the pseudogene term does not make sense, we follow the nomenclature suggested (Cheetham, Faulkner, and Dinger 2020) and only use the term pseudogene alone for sequences that look like genes and are not expressed at all under natural conditions. Consequently, due to limited knowledge of most microorganisms’ natural lifecycles, it is difficult to definitively classify a gene as a vestigial gene or a pseudogene. However, if a gene is rarely expressed under natural evolutionary relevant circumstances and accumulates mutations over time rendering it non-functional, it may eventually become a pseudogene. Several genes close to the *MoISW2* binding sites were not expressed at all in any of the 54 downloaded RNAseq datasets (**Fig. 6B** and **Supplementary Data 4)** indicating that these genes could be or become vestigial genes or pseudogenes without biological roles, thus we call them potential pseudogenes following the definition for the pseudogene concept recently suggested (Cheetham et al. 2020). Our observation that these genes are or have become potential pseudogenes during evolution is exciting since some of these genes close to the MoIsw2 binding sites are annotated as avirulence effector genes (**see Table 1**).

### DNA binding genes

Since 65 genes were classified as DNA-binding and were among the enriched genes upregulated in *ΔMoisw2* (**Fig. 3C, Supplemental Data 13**) these could be conventional TFs, suppressors or other DNA binding genes needed when interacting with the biotic and abiotic environment potentially of high importance during pathogenesis.

The 12 DNA-binding genes most suppressed by MoIsw2 activity are mainly involved in DNA repair (**Table 2**) needed for DNA synthesis and growth. Among the most regulated genes were two TFs and several helicases as well as other genes needed for DNA repair during and after DNA synthesis. A fungal-specific gene involved in (sexual) sporulation is also in this list of 12 DNA-binding genes most suppressed by the MoIsw2 activity.

**Table 2.** Annotation of the 65 DNA binding genes identified in the analysis shown in **Fig. 3C**. These 65 genes are the most upregulated in *ΔMoISW2* compared to the expression in the background Ku80-strain. Annotation from NCBI. Note: these are the genes that are most repressed by the MoIsw2 activity.

|  |  |
| --- | --- |
| MGG_02762 | MGG_02762. ATP-dependent RNA helicase DED1; Belongs to the DEAD box helicase family |
| MGG_06470 | MGG_06470, DNA repair helicase RAD25 (835 aa). |
| MGG_05948 | MGG_05948 zinc knuckle domain-containing protein. This domain is a zinc-binding domain of the form CxxCxxxGHxxx C from various species. It is found in the MPE1 protein from *Saccharomyces cerevisiae* which is a component of the cleavage and polyadenylation factor (CPF) complex important for polyadenylation-dependent pre-mRNA 3'-end formation |
| MGG_01990 | b-ZIP transcription factor IDI-4 (induces autophagic cell death in the fungus *Podospora*) |
| MGG_00868 | Global Transactivator; Superfamily II DNA or RNA helicase, SNF2 family [Transcription, Replication, recombination and repair]; |
| MGG_02429 | KOG4062 6-O-methylguanine-DNA methyltransferase MGMT/MGT1, involved in DNA repair Replication, recombination and repair https://jgi-myco-web-4.jgi.doe.gov/annotator/servlet/jgi.annotation.Annotation?pDb=Lasov1&pStateVar=View&pProteinId=680852&pViewType=protein |
| MGG_07015 | DNA repair protein Rad7 |
| MGG_11518 | G/U mismatch-specific uracil DNA glycosylase |
| MGG_04428 | Zinc finger transcription factor ace1 |
| MGG_07118 | sporulation-specific protein 5 (needed for sexual spore formation?) |
| MGG_05995 | MGG_05995, *Magnaporthe oryzae* 70-15 hypothetical protein (244 aa), translin family protein, Translin family (PF01997). If Translin, here is a review about that https://link.springer.com/article/10.1007/s12038-019-9947-6. |
| MGG_04429 | ATP-dependent DNA helicase MPH1; ATP-dependent DNA helicase is involved in DNA damage repair by homologous recombination and genome maintenance. |

## Functional classification of genes potentially affected by the MoIw2 activity

### A. Genes at the MoIsw2 binding sites in DNA sequences

We have above shown that genes under *MoISW2* control that are more expressed in the background *Ku80* strain compared to the *ΔMoisw2* strain are enriched for gene classes associated with secondary metabolism as well as biomass growth (anabolism) while the genes that are downregulated in the mutant are connected to aerobic metabolism and stress (**Fig. 3C**).

Using the *MoISW2* ChIP-seq data, we investigated the FunCat classification of all genes adjacent to or in ChIP sequence hits to find out what types of genes are overrepresented and which are depleted. First, we removed double ChIP peak sequence hits close to the same gene so as not to count the same gene twice. We then investigated the list of genes that are closest to the found MoIsw2 binding sites. These genes should be targeted by MoIsw2 and might be under positive or negative control from MoIsw2. According to the previous analysis, they should not belong to growth (anabolism) but be genes active when oxygen is consumed, and the substrates oxidized (catabolism) (Klevecz et al. 2004; Machné and Murray 2012). In addition, the binding sequences are located close to avirulence genes and retrotransposons, so their regulation can shift depending on retrotransposon transpositions. Note that since ChIP-seq only shows potential binding to DNA *in vitro,* we cannot say if these genes are up or downregulated; only that a regulation influenced by MoIsw2 is possible.

Genes with ChIP binding hits closest to the MoIsw2 palindromic DNA binding motif site are mainly enriched for genes involved in secondary metabolism and other gene classes important for biotic interactions (**Fig. 7A)**. In contrast, categories of genes for biomass growth and housekeeping (anabolism) are depleted (**Fig. 7B**).

Many other genes within ChIP-seq sequences could be affected. These genes are many more than those closest to the MoIsw2 palindromic DNA motif2 sites. For these (**Fig. 7C**)., we find more gene classes including detoxification and signalling (cyclic nucleotide-binding), disease, and defence, and these are all important gene classes involved in biotic and abiotic interactions with the environments. Depleted are gene classes characteristic for anabolism (biomass growth) (**Fig. 7D**).

Finally, the genes in the ChIP-seq data but not with motif2 hits (not containing the MoIsw2 palindromic DNA binding motif 2 sites) are again genes enriched for secondary genes and again genes involved in abiotic and biotic interactions (**Fig. 7E**). Genes needed for biomass growth are depleted (**Fig. 7F**).

### Functional classification of 500 genes showing the highest VarR and DelR values

As we have seen above (**Fig. 7**) the genes close to MoIisw2 binding motif123 are in general more variable expressed DNA regions between RNAsec plant infection experiments (VarR). These regions number the most TEs and core secondary metabolite genes. To get more information on functional classes regulated we used FungiFun again and did a functional classification of the 500 most variable genes (VarR) in the downloaded data and compared that with the genes most affected by the *MoISW2* deletion (**Fig. 9, Supplemental Data 12**). Few of these genes are probably directly regulated by MoIsw2 but the access to them for gene regulation should be affected. Furthermore, similarities and differences between VarR and DelR should point to regulations biased for genes important for plant interactions or plate growth respectively.

Genes categories needed to cope with the abiotic and biotic environment are enriched among the 500 with the highest VarR **and** DelR values indicating that both the lab and inside plants’ similar gene functions are in principle affected by the MoIsw2 activities (**Fig 9A**). Depleted functional gene classes are growth-related genes (anabolic) and some stress-related genes (**Fig 9B**).

Genes categories needed to cope with the abiotic and biotic environment are again enriched among the VarR 500 most differentially regulated gene classes (**Fig. 9C**). Depleted are again growth-related genes (anabolic) and stress-related gene classes (**Fig. 9D**).

The fungal biomass had recently been shifted from stationary (agar plugs) with the nuclei in various states of growth to a liquid medium for 48h triggering vigorous growth. Thus, variations in the genes affected by the deletion (DelR) are expected in genes for how fast the biomass reacts to the new situation of plenty of nutrients also for genes not affected by MoIsw2 activities. That is probably why unfolded protein response genes are among the enriched genes most affected by the deletion in DelR (**Fig. 9E**), But in general, the functional gene classes among the most regulated VarR alone and most deletion-affected DelR alone are similar.

The specifically differential regulation during plant infection (VarR-specific genes) are genes that are expected to be genes classes responding in fungal-innate immunity (Ipcho et al. 2016) (reacting to plants in this case) (**Fig. 10A-B**) and are further discussed in the discussion section. The DelR-specific enriched and depleted gene classes (**Fig. 10C-D**) indicate gene classes important to regulate during fast aerobic growth of hyphal biomass and nutrient uptake from the environment *in vitro*.

**Figure 8.**
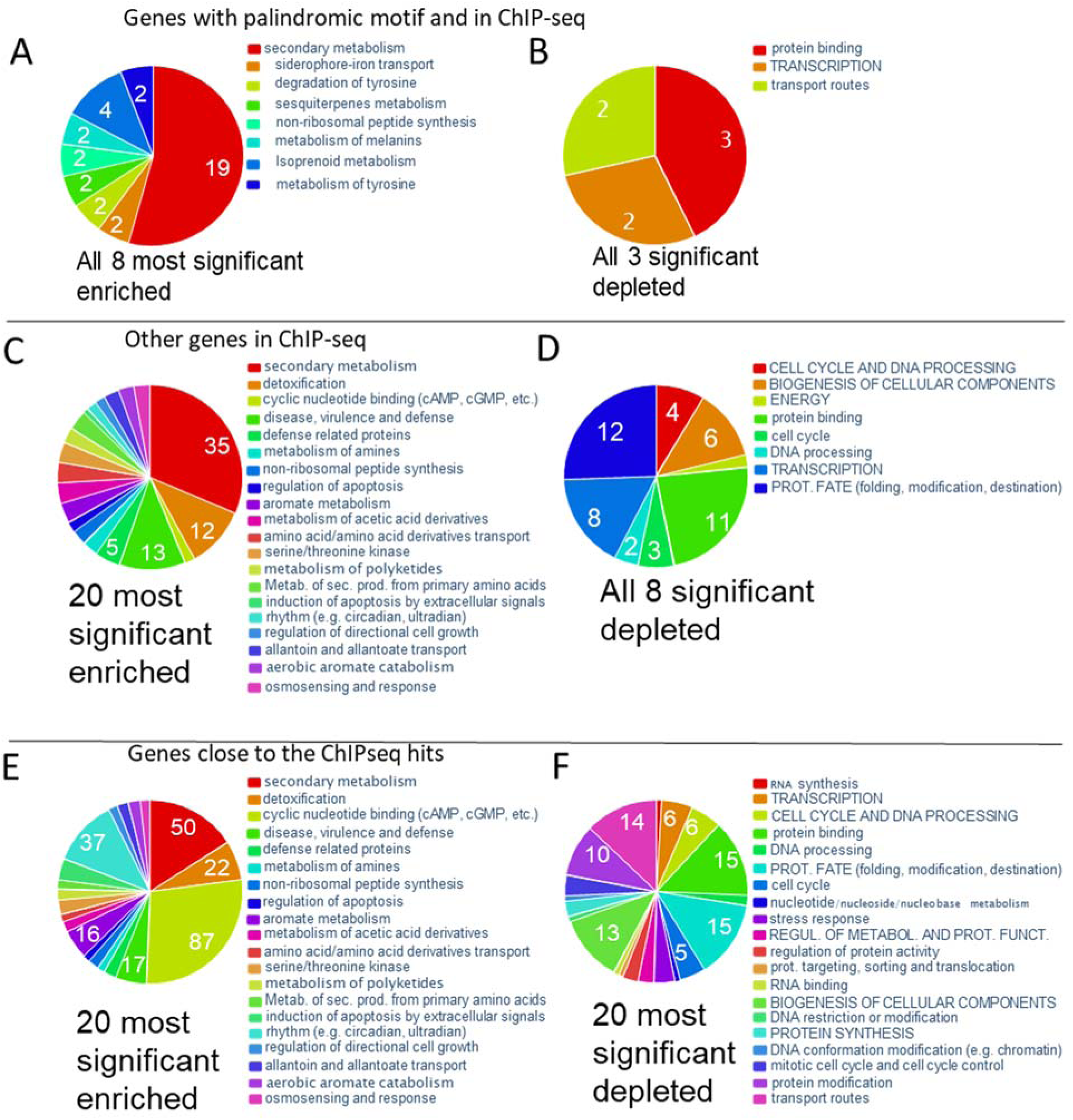
FunCat classification of genes at or close to the MoIsw2 DNA binding sites. **(A)** Significantly enriched and depleted classes of genes in the MoIsw2 ChIP-seq binding assay that also has the MoIsw2 DNA-binding palindromic motif in their upstream region. **(B)** Significantly enriched and depleted classes of all genes with ChIP-seq sequences with MoIsw2 binding to their upstream regulatory DNA sequences. **(C)** Significantly enriched and depleted classes of genes closest to intergenic hits for MoIsw2 binding, excluding the palindromic motif hits.

**Figure 9.**
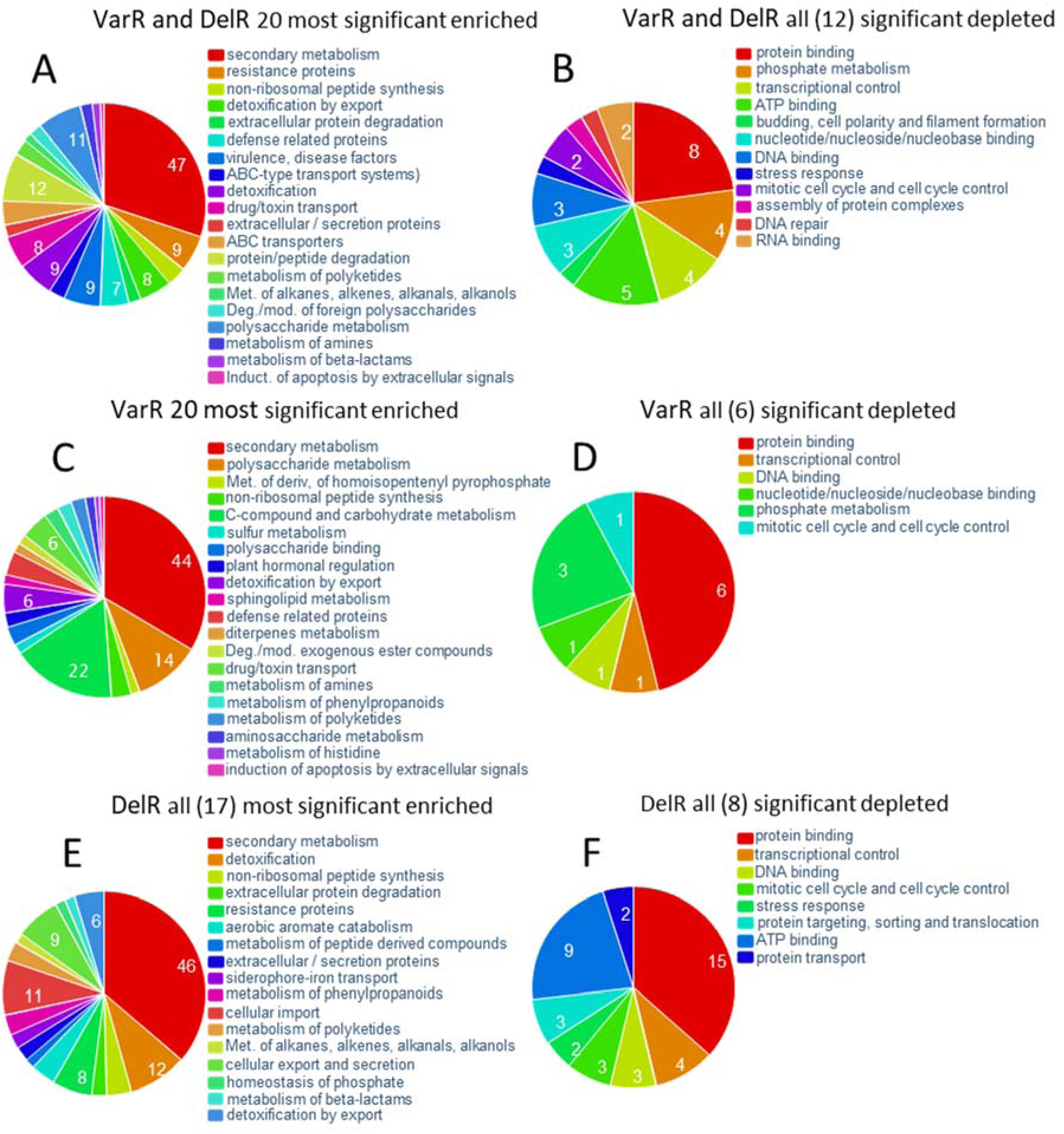
FunCat classification of 500 genes with the most variable gene expression between experiments. (**A**) VarR and DelR enriched. (**B**) VarR and DelR depleted. (**C**) VarR alone Enriched. (**D**) VarR alone depleted. (**E**) DelR alone enriched. (**F**) DelR alone depleted.

**Figure 10.**
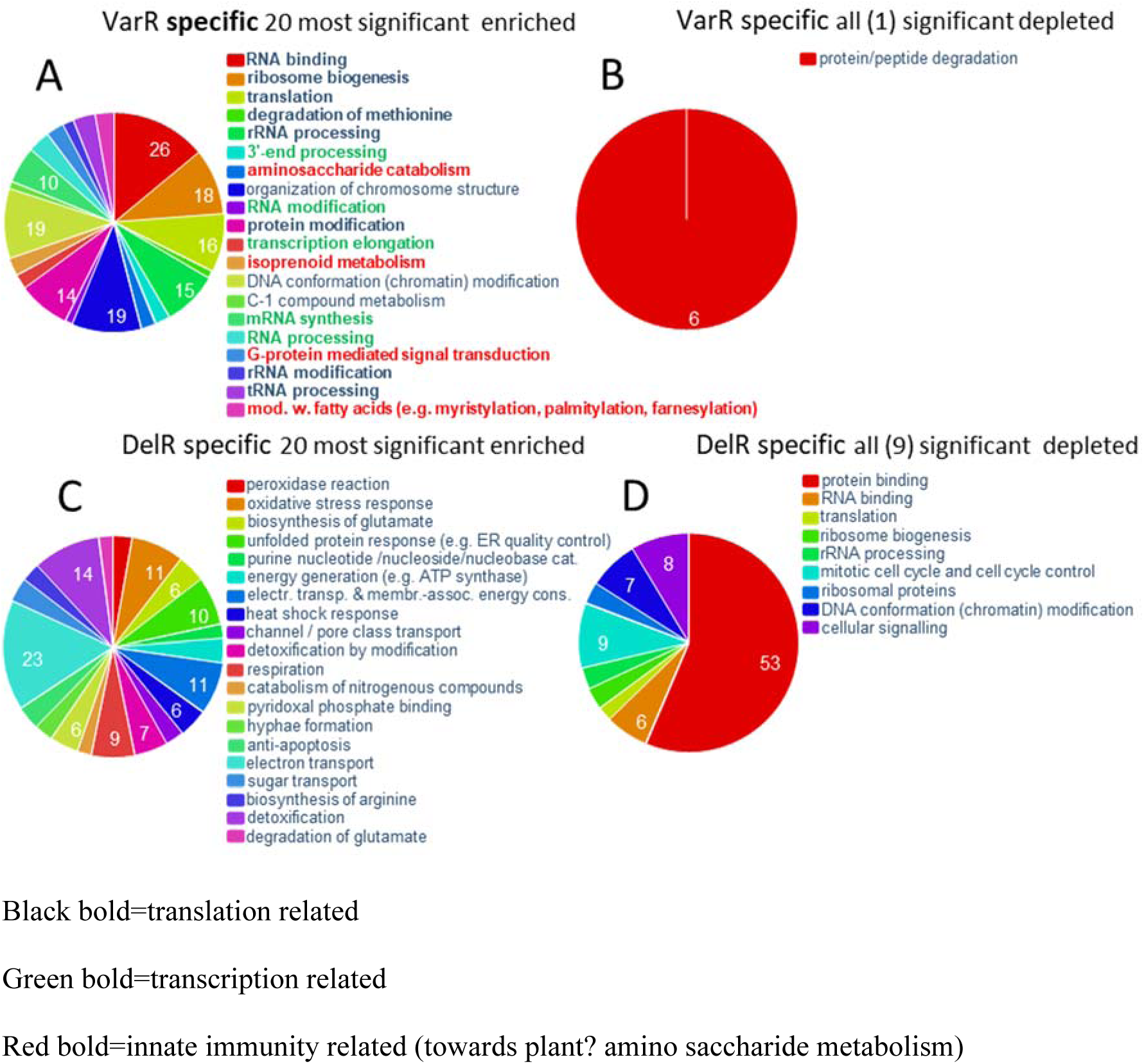
Differences between 500 VarR-specific and DelR-**specific** functional gene classes illustrate the difference in conditions between *in planta* experiments and *in vitro* experiments. (**A**) VarR-specific significantly enriched functional gene classes specifically responding *in planta*. Colour-marked texts are the functional gene classes of special interest for plant pathogenicity (see text). (**B**) VarR-specific significantly depleted functional gene classes specifically responding *in planta*. (**C**) DelR-specific significantly enriched functional gene classes specifically responding *in vitro* (**D**) DelR-specific significantly depleted functional gene classes specifically responding *in vitro*.

## Discussion

### Deletion effects on phenotypes, and regulation (DelR) compared with variability in regulation between RNAseq experiments (VarR)

When *MoISW2* is deleted, the general growth of the fungus (**Fig. 1**) was negatively affected, as was the sensitivity to SDS and NaCl. Conidiation was completely abolished as was also pathogenicity, and the MoIsw2-GFP fusion accumulates in the nucleus as expected for a nuclear localized protein. The strong effect on infection and the fact that Isw2 proteins are known to affect the regulation of many genes in genomes through local chromosomes around several DNA binding sites (Fazzio et al. 2005) indicates that *MoISW2* could be a “master regulator” for regulating the many fungal defences against the plant defences. This is because these and virulence factors (for example secondary metabolism) are especially upregulated in the biotrophy/necrotrophy transition, and during the necrotrophic stage (Andrew et al. 2012; Kou et al. 2019) when membrane effects on the pathogens are especially prominent and the fungus have to deal with plant ROS defences.

We found that potential effector genes like avirulence genes, secreted proteins, and core secondary metabolite genes are located in the DNA regions that appear to be under MoIsw2 control. The protein appears to have the role of a master regulator of TF and repressor gene access to genes necessary for the fungus reacting with the biotic and abiotic environment (**Fig. 5, 6, 7A**) since these gene types are in regions of the DNA that are differentially regulated in the *ΔMoisw2* compared to the background K80 (DelR). These regions are also the same regions with the highest variability between experiments in a set of RNAseq data from 55 downloaded RNAseq datasets from experiments in planta (**Fig. 5**) (VarR). That is what could be expected for nich determinant genes that are not involved in the basic metabolism.

The overall regulation effect of all genes in the two datasets (VarR and DelR) were correlated as they should be if the regulated regions in the two datasets were the same and strain-specific and not experiment-specific and only dependent on the MoIsw2 activity caused organisation of chromatin in the particular strain. Growth inside a plant is stressful for a plant pathogen. The fungus must compete with the plant for all nutrients during the biotrophic stage without revealing its existence and then trigger plant immunity responses that it has to handle in the necrotrophic stage (Andrew et al. 2012; Chowdhury et al. 2017; Rajarammohan 2021). When shifting to the necrotrophic stage the pathogenic fungus must thus both overcome plant defences and nutrient limitations since when spreading inside the plant, nutrient limitations might occur (Josefsen et al. 2012). In support of a strain-specific “master regulator” function for MoIsw2 regulating responses to environmental conditions, we find that many fungal defence and virulence-related functional gene classes are regulated and positioned in the variably regulated chromosomal regions that appear to be under MoIsw2 activity control (**Fig. 8 and 9**). There is in addition a difference in the regulation of avirulence genes between *M. oryzae* strains and these genes are situated close to the MoIsw2 binding sites in the DNA giving the strain specificity of the MoIsw2 as master regulator further support (**Table 1**).

## Regulation of chromatin condensation by nucleosome positioning

### MoIsw2 is a true Isw2

Direct interactions with His4 could not be detected by a yeast two-hybrid assay. The yeast Isw1 and Isw2 interactions with His4 are transient (approximately 200 and 80 bindings per minute, respectively) and need a continuous ATP supply (Tsukiyama et al. 1999). Similarly, the interaction in human cells has been measured in vivo using advanced microscopy and spectroscopy techniques and found to be transient (10-150ms)(Erdel et al. 2010; Erdel and Rippe 2011). Thus, our attempts to detect an interaction between MoIsw2 and MoHis4 using the yeast two-hybrid method were in vain. On the other hand, the expression of MoIsw2 and the expression of orthologues to genes in *M. oryzae,* that an Isw2 protein are known to physically interact and form a functional Isw2 protein complex (Donovan et al. 2021), are correlated (**Fig. 2**). That indicates that MoIsw2 is a true Isw2 with a conserved Isw2 protein function and it has a conserved order of domains in the protein (**Fig. 1A**)

### Regulation of MoIsw2 activity and regulation by *MoISW2* expression

Isw1 translocates nucleosomes away from other nucleosomes, while Isw2 translocates towards the centre of DNA pieces and other nucleosomes *in vitro,*with the nucleosomes bound to relatively short DNA pieces (Kagalwala et al. 2004; Zofall et al. 2004). It does the same *in vivo* where it was found that Isw2 is mainly involved in the targeted regulation of DNA access close to the Isw2 DNA binding site (Fazzio et al. 2005), as we also find (**Fig. 4**, **Table 1**).

The Isw2 activity is ATP dependent (Whitehouse et al. 2007; Hota and Bartholomew 2011; Dang et al. 2014). Thus the regulation of genes adjacent to the DNA-binding site is likely to be highly dependent on ATP availability and competition for ATP in the cytoplasm-nucleus compartment. ATP generation from fermentation is much faster but less efficient than from oxidative phosphorylation when fast fermentation is possible. In yeast, growth is characterized by DNA replication, mainly without mitochondrial oxidative activity (fast aerobic glycolysis), most probably to protect DNA from mutations (Klevecz et al. 2004). In *M. oryzae,* MoIsw2 could thus be an ATP-regulated switch (Machné and Murray 2012) between fast aerobic glycolysis accompanied by DNA-synthesis growth with quality control, and slower but more efficient oxidative phosphorylation growth with high ATP yield, active oxidative defences, and interaction with the abiotic and biotic environment that makes up the *M. oryzae* ecological niche. In support of this, we found that genes upregulated in the *ΔMoisw2* mutant and thus repressed by MoIsw2 under other more ATP-limited aerobic growth conditions are involved in fast growth and DNA synthesis, DNA quality control, while downregulated genes in the mutant were those involved in oxidative phosphorylation, stress management, and secondary metabolite biosynthesis (**Fig. 3EF** and **Table 2**). As also expected from the known function of Isw2, the genes closest to the two central immobilized nucleosomes at the MoIsw2 DNA-binding sites are downregulated in the background *Ku80* (Donovan et al. 2021), and when *MoISW2* is deleted, these genes are upregulated (**Fig. 3A**).

### Regulation by MoIsw2 is dependent on distance from the MoIsw2 DNA binding sites

Gene regulation affected in the *ΔMoisw2* mutant strain compared to the background Ku80 strain (DelR) as well as the variation between RNAseq experiments (VarR) are dependent on the distance from the MoIsw2 DNA binding motif sites (**Fig. 5**). That agrees with that regulation is a consequence of the MoIsw2 positioning of the closest nucleosomes giving more sliding space to surrounding nucleosomes (Donovan et al. 2021) and consequently increased DNA access. Genes affected by the MoIsw2 activities and being physically close to the palindromic binding sites in the DNA are for example genes for secreted protein while core secondary metabolites while typical housekeeping genes like genes encoding ribosomal proteins are much further away (**Fig. 6**). In support of this we found that the 500 most regulated genes (High VarR and/or DelR) with most variable gene expression were significantly enriched for functional gene classes needed to react to the biotic and abiotic environment and depleted for typical housekeeping genes (**Fig. 9**). Interestingly, and in line with this interpretation, the 500 genes specifically upregulated in VarR (during plant pathogenesis) and in DelR (growth in vitro) are genes that reflect the two different biotic/abiotic environments for the fungus. The functional gene classes specially enriched in VarR (especially the ones in red text) are those that are expected for fungal innate immunity (Ipcho et al. 2016) and protection against plant innate immunity like the removal of fungal cell wall derived aminosaccharides (chitin oligomers) that triggers plant immunity (Nürnberger et al. 2004) (**Fig. 10A**).

### Conserved and variable DNA

Our observations suggest that the *M. oryzae* genome is organized into regions with constant expression and variable expression depending on the environment. The regions with variable expression are close to MoIsw2 DNA binding sites and contain genes important for the interaction with the environment (during biotic and abiotic stresses) while regions with less variable expression contain housekeeping genes. Such a division of the fungal genome into regions of housekeeping genes and niche-determinant faster-evolving genes is consistent with previous findings in the relatively closely related *Fusarium graminearu*m (Zhao et al. 2014) but also in *V. dahliae* (Faino et al. 2016).

Palindromic sequences are characteristic of retrotransposons that are mainly positioned close to stress-related genes and avirulence genes in the genomes of fungal plant pathogens. Fungal avirulence genes are plant-pathogen effectors that trigger different plant immunity responses depending on plant variety and the basis for plant cultivar-specific resistance to specific pathogen strains (the gene-for-gene relationship) (Bourras et al. 2016) and we find them close to MoIsw2 binding motifs (**Table 1**, **Fig. 5**). The *MoISW2* DNA binding sequences (the ChIPseq sequences) also contained palindromic DNA motifs characteristic of the identified TEs (Bao et al. 2017) (**Fig. 4**). A palindromic DNA motif gives genetic instability to the genome at the site of the motif (Ganapathiraju et al. 2020; Svetec Miklenić and Svetec 2021). Stress, virulence, and stress-related genes are in *V. dahliae* (Faino et al. 2016) as well as in *M. oryzae,* are also linked to transposable elements (Yoshida et al. 2016) and we see this in our study as well (**Fig. 6, 7A, 8-10**).

Different mutation rates or mutation bias of genes have recently been described as a new concept for the plant *Arabidopsis thaliana* (Monroe et al. 2022). Genes with higher mutation rates are involved in biotic and abiotic interactions. The gene found with the highest mutation rate was a gene responding to chitin, which is present in symbiotic (pathogenic and nonpathogenic) fungi and insects (Monroe et al. 2022). Transposable element activity seems important in fungal speciation (Faino et al. 2016) and tends to accumulate in genomes and expand the genome with repetitive sequences (Fijarczyk et al. 2022; Feurtey et al. 2023) unless there is some evolutionary constraint against such expansion (Kremer et al. 2020; Fijarczyk et al. 2022). Phosphorous availability is a possible constraint (Ågren et al. 2012) since a considerable amount of the cell phosphorous, especially, is bound up in DNA and not available for other purposes (Berdalet et al. 1994). Fungal plant parasites can get phosphorous directly from their plant hosts by degrading the plant tissue and are not so restricted by limited resources in the soil (Ågren et al. 2012). Thus plant parasitic fungi should be expected to have more retrotransposons to adapt better to plant resistance and thus have relatively large genomes. That is precisely what they have (Fijarczyk et al. 2022; Feurtey et al. 2023). Endophytic fungi that have to coexist with their host in their life cycle, and thus have to compete with the host for phosphorous, should, in consequence, have smaller genomes, as they also seem to have (Fijarczyk et al. 2022).

Our results show changes in occurrence and regulation of the avirulence genes between two *M. oryzae* strains with different pathogenicity to different rice cultivars (**Table 1**). In addition, we did not detect any expression for some putative avirulence genes close to the MoIsw2 DNA binding sites, suggesting that those genes may be potential pseudogenes in these strains. That might indicate a retrotransposon-aided genetic evolution of inherited epigenetic changes to regulation in reactions to the environment (resistance of the plant cultivar) as has been indicated for *V. dahliae* (Faino et al. 2016). Such regulation can become genetically fixed, first through loss of possible regulation (making downregulated genes into pseudogenes), and much later complete gene loss or change of the potential pseudogenes so they cannot be easily recognized as genes, is a mechanism that could result in a “biased” faster evolution (Chadha and Sharma 2014). Most of the potential pseudogenes (not expressed under any conditions reported in this paper) were close to the MoIsw2 DNA binding sites with palindromic motifs supporting this hypothesis for a mechanism involving MoIsw2 activity in creating pseudogenes and complete removal of genes by making them unrecognizable as genes through mutations. A prerequisite for this to happen is that the ecological niche does not change as fast as it can for a plant pathogen because of efforts of resistance breeding of susceptible plant cultivars. These efforts can then reactivate the silenced genes by TE repositioning to chromosome areas with changed MoIsw2 activity.

Our results further indicate that *MoISW2* regulates a switch between fast glycolytic growth with fast DNA synthesis and slower but nutrient-efficient aerobic stress-characterized growth with less DNA synthesis, coping with abiotic and biotic stresses or, in other words, niche fitness. The loss of *ISW2* function in a unicellular Eukaryote or the early stages of fungal growth from a spore should make the fungus unfit for survival in a natural environment and become more dependent on aerobic fermentation for ATP synthesis that is not substrate efficient although generates ATP at a very high rate although inefficient (Pfeiffer and Morley 2014; Desousa et al. 2023). However, a limitation for eukaryotic cells to only aerobic fermentation generally leads to uncontrolled cell growth and oncogenesis in a long-lived multicellular organism (Desousa et al. 2023), as ATP and fermentable nutrients can be received from surrounding cells and other tissues through the blood circulation. Thus, mutations of the genes *SMARKA2* or *SMARKA4* that encode Brm and Brg1 proteins that are orthologues of fungal Swi2/Snf2 can cause cancer (Reisman et al. 2009; Mittal and Roberts 2020; Li et al. 2021).

On the other hand, a general shift to aerobic glycolysis and a fast upregulation (within a few hours) of innate immunity-related genes and their ATP-dependent translation is associated with tissue inflammation. Such fast de-novo protein synthesis is characteristic of fast defence against microbes needing rapid translational responses and has been shown for *Fusarium graminearum* but is general for eukaryotes (Ipcho et al. 2016). Such fast upregulated plant innate immune responses can be inhibited by fungal-derived immune system inhibitors acting on translation like trichothecenes the fungus (Toyotome and Kamei 2021), most likely to attenuate immune responses in the plants it infects. In line with a fungal innate immunity scenario (Ipcho et al. 2016), genes involved in RNA synthesis and RNA processing were found to be upregulated in the *ΔMoisw2* mutant (**Fig. 3C**).

### Is retrotransposon shifting *MoISW2* binding creating and directing the mutation bias by adapting the fungus to new challenges?

*MoISW2* is likely an important player in a mechanism aiding both mutation bias and adaptation to new circumstances whose combined activities lead to two-speed evolution of the fungal genome (Faino et al. 2016) a slow random evolution and a fast more directed by adaptations. The faster speed evolution can be named natural adaptation-directed fast evolution (NADFE), a new concept that is similar to artificial adaptation-directed fast evolution that can be performed in a laboratory (Packer and Liu 2015). The suggested mechanism is the following: As transposon DNA containing MoIsw2 palindromic DNA binding sites seems necessary for MoIsw2 activity, stress-activated transposon activity, known for creating genomic instability, can together with the MoIsw2 activity create a new adaptive regulatory landscape (Chadha and Sharma 2014) that can be stabilized relatively quickly and lead to mutation bias (Monroe et al. 2022). In addition, the Isw2 stabilized expression landscape further stabilized by later mutations could be the mechanism sought after for a Lamarckian-Darwinian synthesis. This mechanism can explain the fast evolution of new traits requiring many combined mutations that cannot be reconciled within Darwinian strict random evolution creating variation together with sexual recombination and natural selection to cause both the observed direction and the needed speed of evolution of specific traits when new niches become available, or there are drastic niche changes (Bard 2011). The chance of multiple mutations that only together can become positive is theoretically much more likely to happen with NADFE first creating a mutation bias (Monroe et al. 2022).

### Perspectives for Future Research

*Evolution of the plant-pathogen interaction:* The effect of exposing *M. oryzae* strains to different MoIsw2 activities and retrotransposon activities on the rate of genetic adaptation to a shift in rice cultivar is a possible future topic that could be exploited to understand how the pathogen adapts to different resistant cultivars and break plant resistance.

*Agrochemicals:* Searching for potential agrochemicals to manipulate MoIsw2 activities on His4 and DNA-binding is tempting since *MoISW2* is crucial for plant pathogenicity. Though MoIsw2’s close resemblance to mammalian proteins with the same function indicates that such chemicals are likely negatively affecting orthologue proteins in mammals and might be cancerogenic to humans since even small mutations in *ISW2* orthologues in mammals can cause cancer (Reisman et al. 2009; Mittal and Roberts 2020). ***Histon4:*** Finally, we found 2 *MoHIS4* genes with large differences in DNA sequences (gene duplication many generations ago) predicted to encode for identical proteins but differently regulated and it seems like *M. oryzae* needs both genes (**Fig. 2**). This fact is of interest since His4 proteins in nucleosomes is what MoIsw2 is supposed to interact with (Tsukiyama et al. 1999), and *HIS4*s are examples of genes with the highest degree of purifying evolution (Piontkivska et al. 2002). We have thus started new research into the roles and regulation of these two *MoHIS4* genes and how their regulations shift with growth conditions. Their predicted identical amino acid sequences indicate that they are evolutionarily constrained to differ only by synonymous mutations.

*Gene expression variability:* The most interesting genes for biotic and abiotic interactions are also the genes with the highest variability between experiments (VarR) creating problems when proving the effects gene deletions. The common practice of only using 3 replicates for measuring effects on genes (like deletions) is probably not enough and we might need to increase the number of replicates to 5 for the gene expression in the wild type. If we do that we can either statistically prove what type of distribution there is and use that or better use non-parametric methods not assuming the distribution. Strict normal or more correctly lognormal distribution (negative expression is not possible!) may also be true mainly for housekeeping genes.

### Consequences of MoIsw2 functions that are positive for the fungus’s ability to stay pathogenic

The MoIsw2 function will result in adaptive silencing of virulence/avirulence genes without gene loss due to mutations of the actual DNA nucleotides. In other words, virulence genes are not quickly lost from the adapted strains but can be “called upon again” if the available hosts change their resistance in the classical gene-for-gene scenario (Flor 1956). 2. Adaptive silencing of virulence genes is often found close to transposable elements (Menardo et al. 2017) and that will lead to biased fast gene evolution of especially genes involved in biotic interaction (Monroe et al. 2022) and in the gene-for-gene concept and is a possible mechanism behind the speedy appearance of virulent pathogen strains capable of attacking newly developed resistant plant varieties (Palloix et al. 2009; Menardo et al. 2017).

## Materials and Methods

### Fungal strains and media

*Magnaporthe oryzae* B. Couch anamorph of the teleomorph *Pyricularia oryzae* Cavara was used for this research. As background strain, we used Ku80 (generated from the WT strain 70-15) to minimize random integration events when transformed (Villalba et al. 2008). The susceptible Indica rice (cv. CO-39) and barley (cv. Golden Promise) used for the fungal pathogenicity tests were from the seed bank of our laboratory. For both ChIP-seq and RNAseq, the strains used were grown and harvested similarly.

### Knockouts, complementations, and verifications

The *MoISW2* is a *MYB* gene and *MYB* gene deletion vectors were constructed in the plasmid pBS-HYG by inserting 1 kb up– and down-stream fragments of the target gene’s coding region as flanking regions of the HPH (hygromycin phosphotransferase) gene (Li et al. 2012). No less than 2 μg of the deletion vector DNA of the target gene was introduced to Ku80 protoplasts, and transformants were selected for hygromycin resistance to perform gene deletion transformations. Southern blotting was conducted to confirm the correct deletion using the digoxigenin (DIG) high prime DNA labelling and detection starter Kit I

(11745832910 Roche Germany). The *MYB* gene complementation vectors were constructed by cloning the entire length of the target gene with the native promoter region (about 1.5 kb) to the pCB1532 plasmid. When making the complementation vector, GFP was linked to the C-terminal of the target genes to study the sub-cellar localization of Myb proteins. The constructed vector DNA was introduced into the mutation protoplast for gene complementation, and resulting transformants were screened using 50 μg/ml chlorimuron-ethyl to select successful complementation strains. Detailed fungal protoplast preparation and transformation methods have been described previously (Li et al. 2012). All primers needed for the knockout and complementation are listed (**Table S1**). The sub-cellar localization of Myb proteins was observed by confocal microscopy (Nikon A1). GFP and RFP excitation wavelengths were 488 nm and 561 nm, respectively.

### Colony growth and infection phenotype measurements

Vegetative growth was tested by measuring the colony diameter after ten days of growth in 9 cm Petri dishes at 25℃ under 12h-to-12h light and dark periods. Conidia production was evaluated by flooding the 12-day-old colony with double distilled water, filtering out the mycelia with gauze, and counting the conidia using a hemacytometer. The conidiophore induction assay was performed by excising one thin agar block from the fungal colony and then incubating it in a sealed chamber for 24 h with constant light (Li, Yan, et al. 2010). Mycelia appressoria was induced by placing a suspension of mycelial fragments on a hydrophobic surface in a humid environment at 25℃ for 24h. The pathogenicity assay on rice was performed by spraying 5 ml conidial suspension (5 × 10^4^ spores/ml) on 15-day-old plants (Y. Li et al. 2019). The inoculated plants were kept in a sealed chamber with a 90% relative humidity at 25℃ for 24 h before the inoculated plants were removed from the chamber to allow disease symptoms to develop for 4-5 days. The pathogenicity assay on excised barley and rice leaves was performed by cutting a small block from the agar culture of the fungus and placing it on excised leaves for five days in a moist chamber for disease development (Li, Liang, et al. 2010). Sexual reproduction was tested by crossing the tested strain with the sexually compatible strain TH3 on OM plates and then incubating at 19℃ for 30 days with continuous light. The perithecia and clavate asci were photographed in a microscope equipped with a camera (OLYMPUS BX51).

### Biomass production for ChIPseq and RNAseq

For ChIP-seq, the complemented strain, MoIsw2-GFP and the strain Ku80 were used, and for RNAseq *ΔMoisw2* and *Ku80* were used. The strains were grown on Complete medium 2 (CM2) all are quantities L^-^: 20X Mineral salts solution 50mL, 1000X Trace element solution 1mL, 1000X vitamin solution 1mL, D-glucose 10g, peptone 2g, casamino acid 1g, yeast extract 1g, pH 6.5. For agar medium add 15g agar. Supplemented with 20×Mineral salts solution (per 1000mL): NaNO3 120g, KCl 10.4g, MgSO4 7H2O 10.4g, KH2PO4 30.4g and with 1000X Vitamin solution (per 100mL): Biotin 0.01g, Pyridoxin 0.01g, Thiamine 0.01g, Riboflavin 0.01g, PABA (p-aminobenzoic acid) 0.01g, Nicotinic acid 0.01g and 1000X Trace element (per 100mL): ZnSO_4_ 7H_2_O 2.2g, H_3_BO_3_ 1.1g MnCl_2_ 4H_2_O 0.5g, FeSO_4_ 7H_2_O 0.5g CoCl_2_ 6H_2_O 0.17g, CuSO_4_ 5H_2_O, Na_2_MoO_4_ 5H_2_O 0.15g, EDTA 4Na 5g. The plates were incubated for 4-5 days at 28°C with alternating light-dark cycles (12/12). Mycelial disks (15-20) were punched out at the colony’s edge using a 5 mm diameter cork puncher. The disks were transferred to 100 ml CM2 liquid medium and shake cultured for another 48h (160 RPM, Constant temperature culture oscillator, ZHWY-2102C, Shanghai Zhicheng Analytical Instrument Manufacturing Co., Ltd.). The obtained biomass was filtered using Miracloth, frozen in liquid nitrogen, and sent on dry ice to the company performing the RNAseq.

### ChIP-seq and RNA-seq

Both these techniques were carried out by the company Wuhan IGENEBOOK Biotechnology Co., Ltd, China, according to their method descriptions **(Supplemental Company Methods file**). The last steps about finding motifs and enriched gene classes were carried out differently and are described in this paper. These steps not used are marked in the two Supplementary method files.

### Method for calculating VarR and DelR in the RNAseq data

We calculated the variability in gene expression for each gene for all the downloaded experimental datasets in the following way. First, we checked that the quality of the data was ok. We have previously found for our own transcriptomic data that a dataset is ok if expressions of genes are rank-ordered then the log log expression of log expression level against the log-rank order should be a smooth downward slightly curved line without big gaps. Also, to be able to compare different sets of data from different experiments the total expression of all genes summed should be similar for all datasets. Next, we calculated the maximum variation between experiments in the following way.

The maximum expression value and the minimum expression value for each gene across the 55 transcriptomes were found. Next, we calculated the average expression for the same data. Finally, we calculated the ratio of maximum expression minus minimum expression divided by the average expression to arrive at the variation in expression across all experiments for each gene (VarR). In this way any regulation up or down is treated in the same way as access for TFs and repressors to the DNA should be treated equally. Finally comes the problem that some genes had expression values lower than the threshold for detecting them at all (0-values). For these data, we assumed that the expression where at the threshold value instead of 0 and set the expression to that value. In that way, we could include that data without overestimating the VarR for that gene.

The effect of the *MoISW2* mutation (DelR) was estimated in the following way to make it comparable with VarR: For each gene, we calculated the 2Log ratio of the expression in the mutant divided by the expression in the background.

That results in both negative and positive values. Then these values were squared to arrive at only positive values in a similar way as is done with the least square method for curve fitting to arrive at a better estimate of the deviation from the slope. There were very few genes with no detectable expression so no need for handling zero values more than excluding them.

Note that in this paper, we focus on the expression variation of all genes along the chromosomes. Thus, it is the overall pattern that is important and not the values for a single gene. In none of the methods above we used any method for excluding a gene from the analysis based on an arbitrary threshold of 2 times up or down or any other statistically significant threshold for genes. In consequence, the calculated values for a single gene cannot and should not be relied upon even if a general increased regulation of many adjacent genes has reliability.

### Additional software and add-ins for MS Excel

***Addins for MS Excel:*** The **Fisher Exact** add-in for MS Excel was downloaded from http://www.obertfamily.com/software/fisherexact.html. The **Excel solver** was used to fit non-standard equations to data. The Solver is usually part of MS Excel but must be activated in settings.

**Freeware:** We used the freeware PAST A simple-to-use but powerful freeware program is available from the University of Oslo, Natural History Museum https://www.nhm.uio.no/english/research/infrastructure/past/ (Hammer et al. 2001) version 4.08 (released November 2021). We mainly used the software for Reduced Major Axis (RMA) regression analysis to handle errors in both the x and y variables. For simplicity, the data was handled and entered in MS Excel and then copy-pasted into PAST for analysis. The resulting plots were exported from PAST as SVG vector graphic files to be later translated into other vector graphic file formats.

## Websites used for analyses and getting necessary additional data

**NCBI**: https://www.ncbi.nlm.nih.gov/ (sequence downloads, blasts, annotations including domain annotations)

**BROAD**: ftp://ftp.broadinstitute.org (download of gene order on supercontigs for the Guy11 strain, 1000 upstream DNA sequences for use in MEME)

**MEME:** https://meme-suite.org/meme/ (use of MEME and FIMO). (Bailey et al. 2015).

See the supplemental file for MEME settings and complete results (Supplemental Data 2). MEME motif 2 was the one with the most hits and was investigated in detail. This motif2 was used for TOMTOM by using the direct link to TOMTOM using the link from the MEME result, for results see the supplemental file (Supplemental Data 3).

**FungiFun2** https://elbe.hki-jena.de/fungifun/fungifun.php (Priebe et al. 2015) Species: *Magnaporthe grisea* 70-15 (synonym for *M. oryzae* 70-15).

Classification ontology: FunCat (contains more relevant categories for fungal plant pathogens than Kegg or GO)

Input IDs: For *M. oryzae* 70-15 the MGG_ codes has to be used.

Advanced settings used: Significance level; 0.05. Significance test; Fisher’s exact test. Test for; Enrichment or Depletion. Adjustment method; No adjustment. Annotation type, Select also indirectly annotated categories.

These settings using “No adjustment” were used since the purpose was not to get super reliable enriched or depleted categories but to compare several FungiFun runs with different Gene ID sets.

## antiSMASH

The complete DNA sequence of *M. oryzae* 70-15 was downloaded from BROAD. That was then submitted to antiSMASH (Blin et al. 2021) fungal version https://fungismash.secondarymetabolites.org/#!/start and run with default parameter settings to find the core genes predicted to be involved in producing secondary metabolites.

## Supporting information

All supplemental Figs and tables

## Acknowledgements

This work was supported by grants from the Fujian Provincial Science and Technology Key Project (2022NZ030014) Fujian Natural Science Foundation project (2022J01125).

---

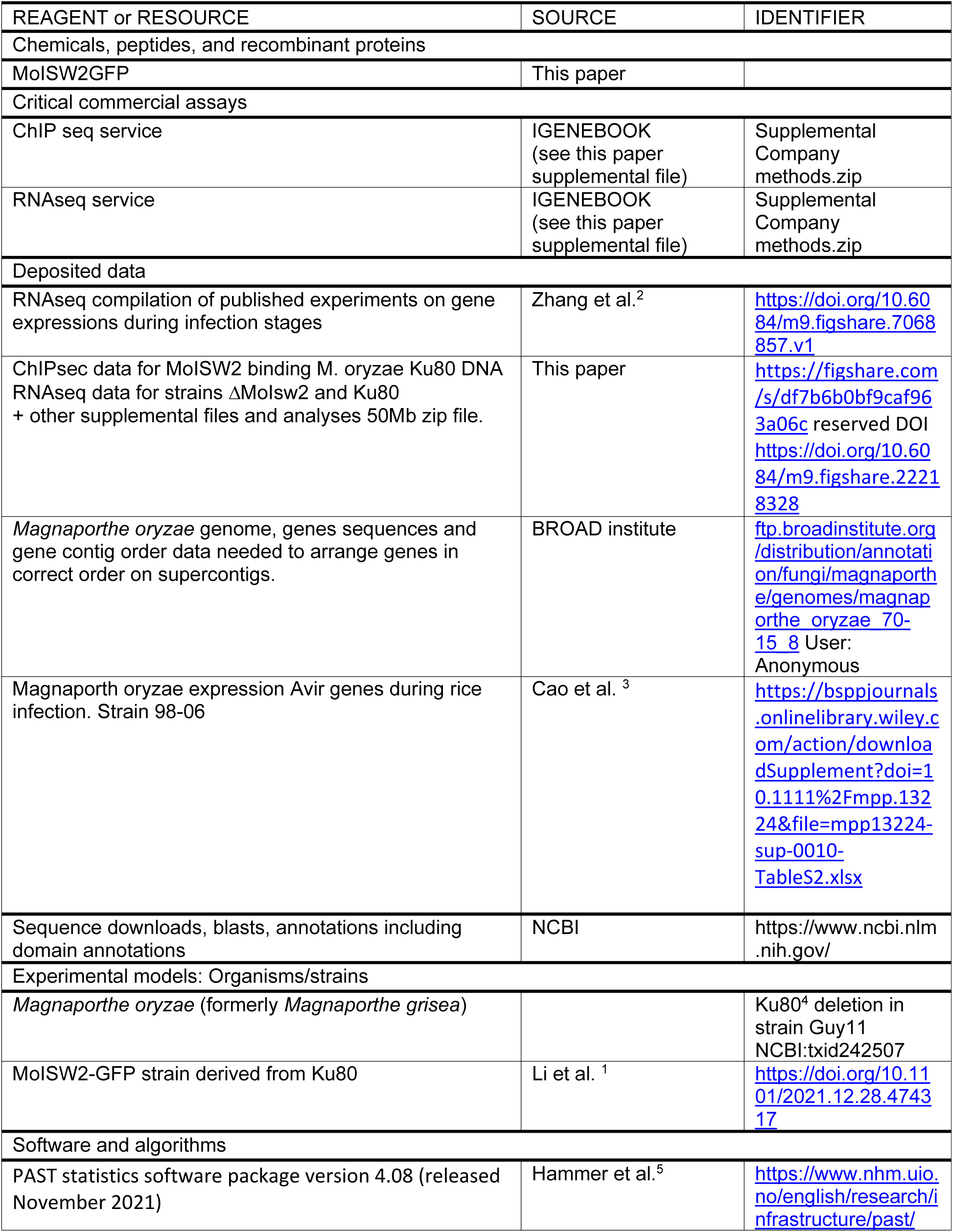
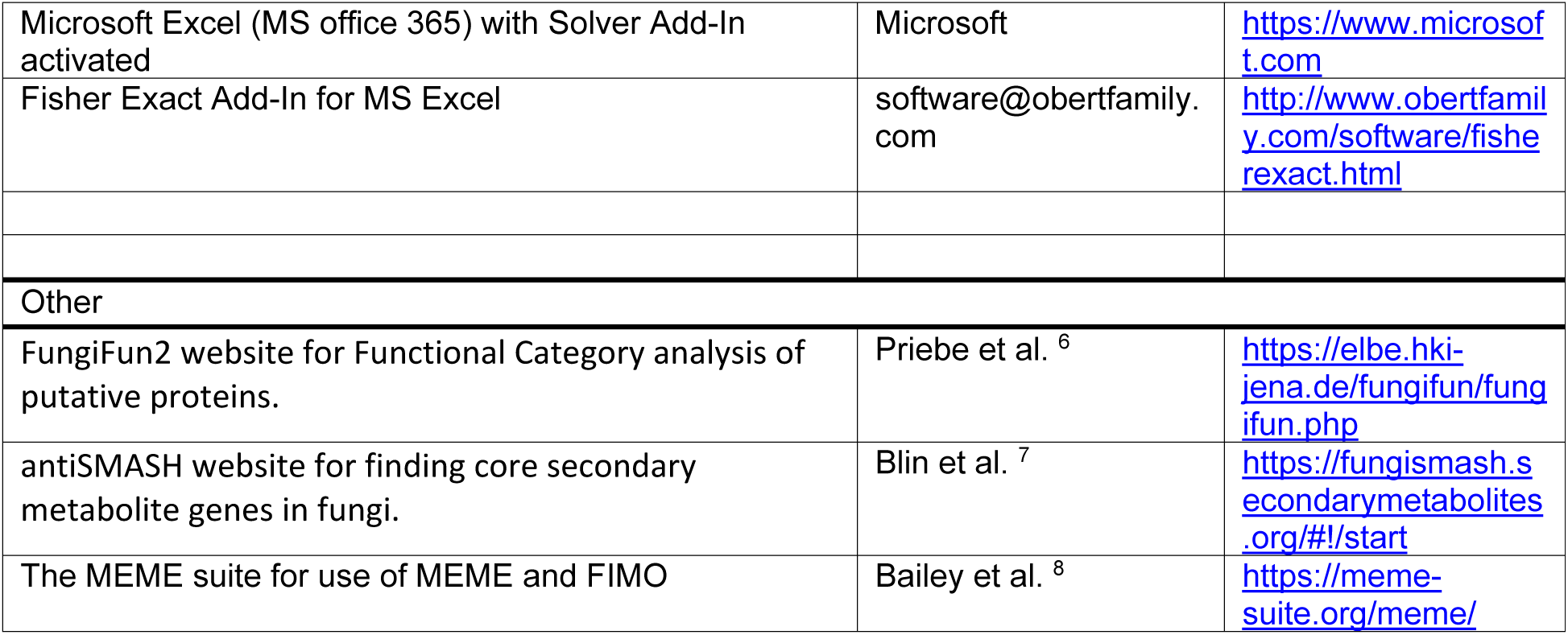
Key resources table.

