## Supplementary material for "Whole genome regulatory effect of *MoISW2* and consequences for the evolution of the rice plant pathogenic fungus *Magnaporthe oryzae*": All supplemental Figs and tables

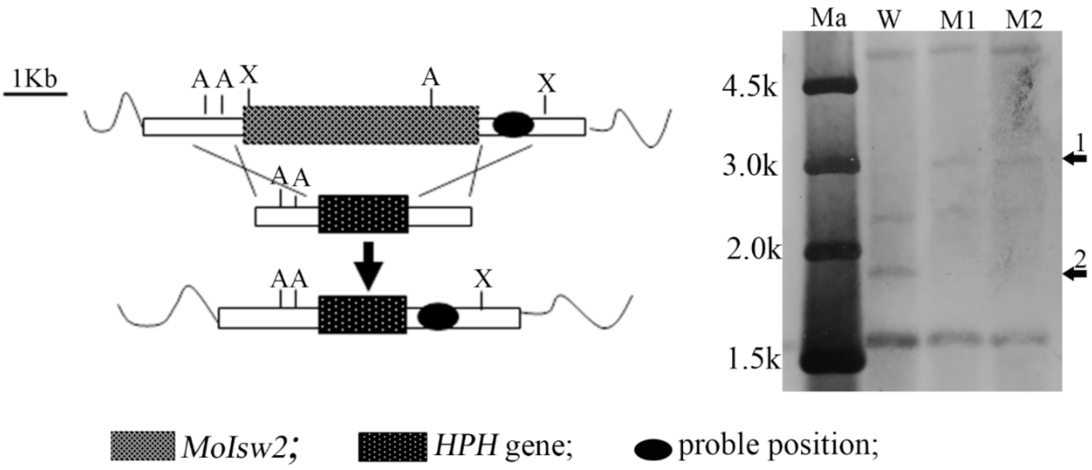


Figure S1. Southern blot confirmation for MoIsw2 mutants. The 1.0 kb downstream of the target gene was used as a probe for blotting. The arrow1 indicates the expected bands that should be present in mutants. The arrow2 indicates the expected bands that should not be present in the mutants, but present in Wild-type strain. W, Wild-type strain; M1 and M2, two mutants of the target gene; Ma, Mark; A, AgeI; X, XhoI;

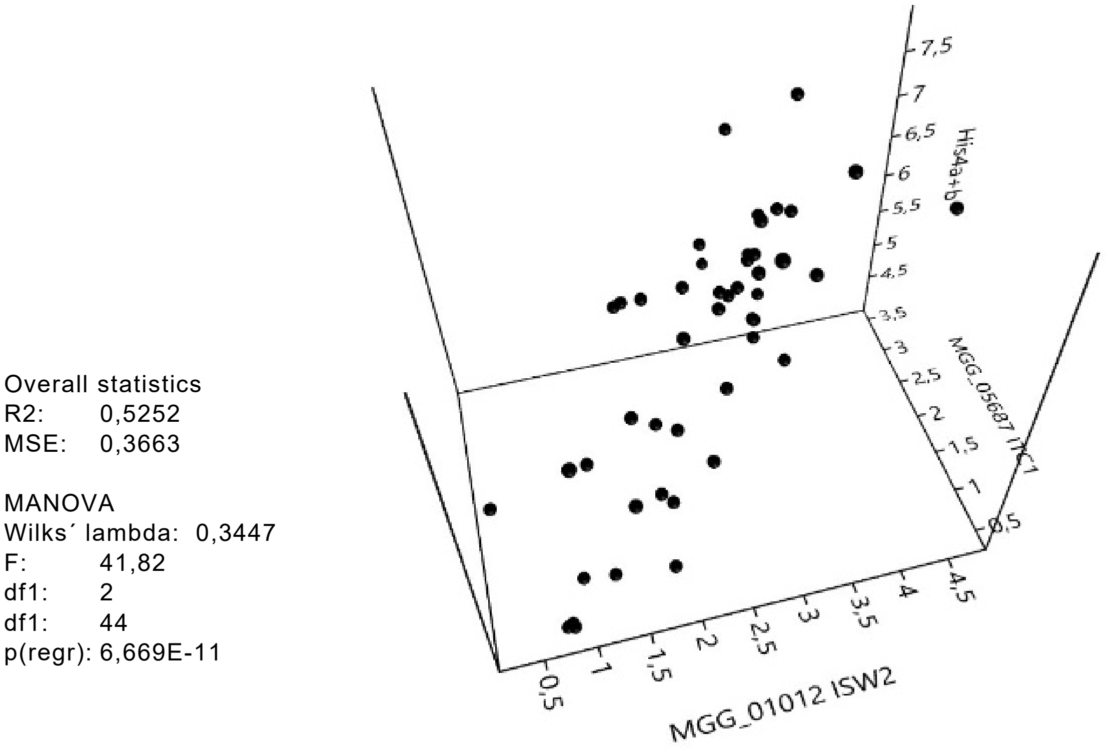


**Figure S2.** 3D plot showing the Log of expression of *MoISW2*, *MoITC1*, and *MoHIS4a+b* that, according to the literature, need to work together to influence nucleosome packing. They are all correlated and form a nice 3d "sausage" as they should if the three genes encode proteins that need to work together. Multiple regression with *MoISW2* as an independent variable, and *MoITC1*, and *MoHIS4a+b* dependent gives a linear relationship (P=6.7E-11).

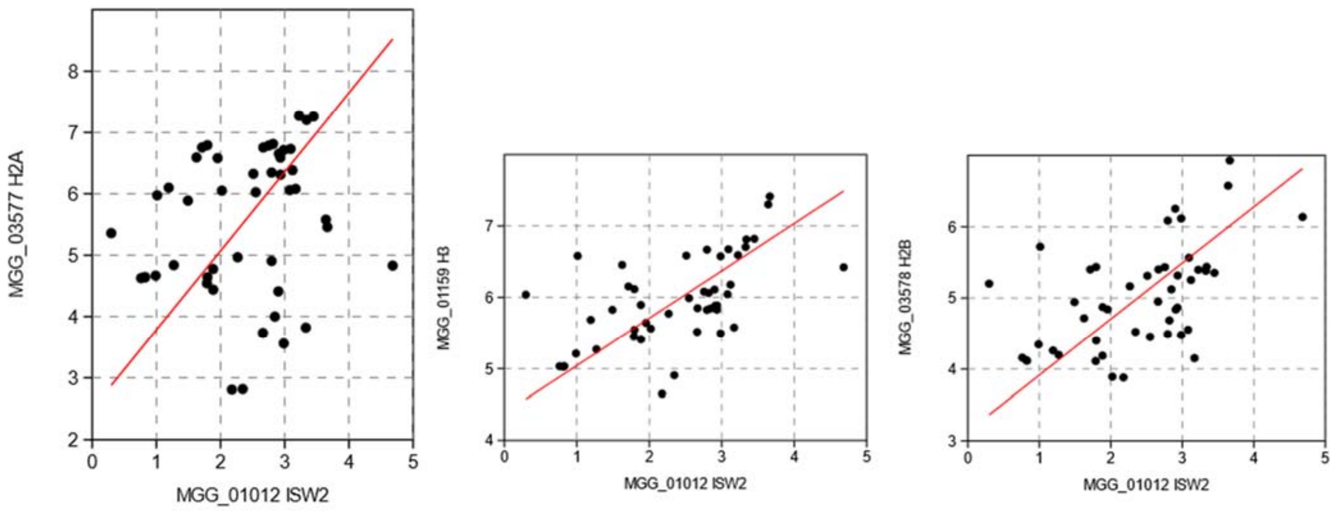


**Figure S3.** Log2 RMA correlations of a putative *MoHIS2a, MoHIS3* and *MoHIS2B* with the expression of *MoISW2* (x-axis) in published RNAseq data at different stages of plant infection. (**A**) *MoHIS2a*, P(uncorr=0.20 (**B**) *MoHIS3*, P(uncorr)=8.94E-5 (**C**) *MoHIS2B*, P(uncorr)=3.56E-5.

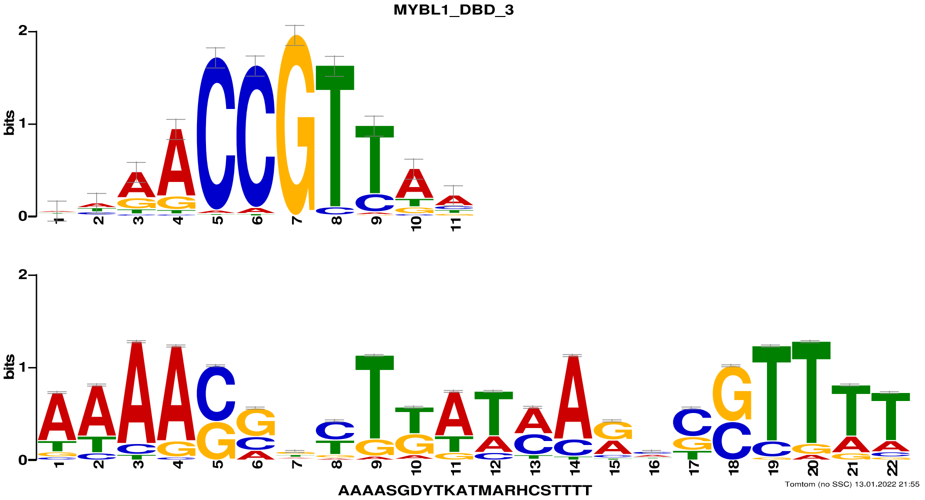


**Figure S4.** Comparison of the motif (Bottom) with the known human Myb protein DNA binding motif (Top) found by a TOMTOM query using the motif in Fig. 2F as the query. The MoIsw2 DNA binding site has a palindromic Myb/SANT-like DNA binding motif with a similarity to the human protein MYBL1.

Correlation LOGDelR vs LOG VarR data

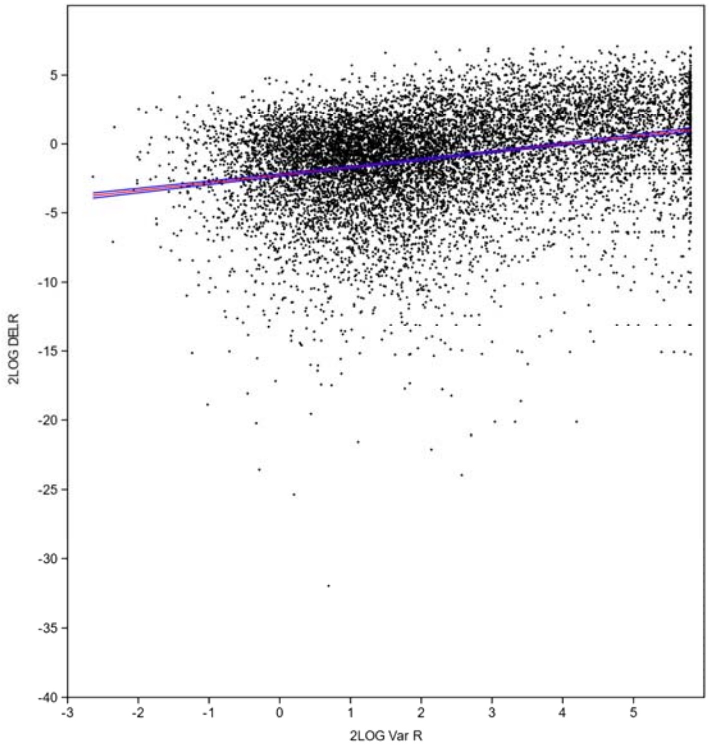


Ordinary Least Squares Regression: 2LOG Var R-2LOG DELR

| | | | | |
|---|---|---|---|---|
| Slope a: | 0.56337 | Std. error a: | 0.018482 | |
| | t: | 30,482 | p (slope): | 5.2724E-196 |
| Intercept b: | -2,2609 | Std. error b: | 0,051775 | |

95% bootstrapped confidence intervals (N=1999):
Slope a: (0.52552, 0.60006)
Intercept b: (-2.3619, -2.1567)

Correlation:
r: 0.27884
r2: 0.077754
t: 30.482
p (uncorr.): 5.2724E-196
Permutation p: 0,0001

The red plot line shows the calculated fit to the data
The blue plot lines show the 95% confidence level lines for the fitted red line.

**Figure S5** Correlation between 2Log DelR vs 2Log VarR data. The VarR responses in planta and the DelR responses in vitro are correlated. Genes with very low VarR variability are good candidates as reference genes for the *in planta* experiments. Genes with low DelR variability are good candidates for the in vitro experiments. The commonly used reference gene that seemed to show very low variability was the actin gene. The technique should be able to be used to select good and stable reference gene candidates for RNAseq for different conditions. It should be able to find genes stable enough that small changes in the expression of target genes like TFs and other regulators and receptors can be detected since thos gene classes can have large biological effects even with minor expression changes.

### Table S1. Primers used in this study

| Name | Sequences | Application |
|---|---|---|
| MGG_01012-UP-F | aagggaacaaaagctggtaccACGGCTTGACGGTTACTTGT | Gene deletion |
| MGG_01012-UP-R | tcagttaacgtcgacaagcttTTGTCGTGAGATTCCCTGGT | Gene deletion |
| MGG_01012-down-F | cgggaaccagttaacctgcagGTTTGGAACTCTTTGATGGG | Gene deletion |
| MGG_01012-down-R | cgctctagaactagtggatccGGGCTACTTTGACTTTATGT | Gene deletion |
| pCB1532-MGG_01012-Pro-F | cgctctagaactagtggatccAGGTGGATGATGTCGATTGCC | Gene complementation and protein localization |
| pCB1532-MGG_01012-EcoR1 | gcccttgctcaccatgaattcTTTCTTCTTGCCCTTGGCC | Gene complementation and protein localization |
| MGG_01012-TZ-F | gactttcgggacaagatgga | Southern blot probe |
| MGG_01012-TZ-R | acacccatcgcgaatagaac | Southern blot probe |
